# LOCAL SPATIAL FLOWS AND PROPAGATIVE ATTRACTORS: A NOVEL “FLOWNECTOME” FRAMEWORK FOR ANALYZING BOLD FMRI DYNAMICS

**DOI:** 10.1101/2024.01.26.576775

**Authors:** Robyn L. Miller, Victor M. Vergara, Erik B. Erhardt, Vince D. Calhoun

## Abstract

Although the analysis of temporal signal fluctuations and co-fluctuations has long been a fixture of blood oxygenation-level dependent (BOLD) functional magnetic resonance imaging (fMRI) research, the role of local directional flows in both signal propagation and healthy functional integration remains almost entirely neglected. We are introducing an extensible framework, based on localized directional signal flows, to capture and analyze spatial signal propagation and propagative attractor patterns in BOLD fMRI. Novel features derived from this approach are validated in a large resting-state fMRI schizophrenia study where they reveal significant relationships between spatially directional flows, propagative attractor patterns and subject diagnostic status. Plausibly, we find that spatial signal inflow to functional regions tends to positively correlate with net gain/loss in the region′s temporal contribution BOLD signal reconstruction. We also find that group ICA (pdGICA) performed on time-varying propagative density maps, which are whole-brain maps of spatial signal inflow to each voxel on successive 30 second windows (ie. propagative attractor maps), produce components that tend to concentrate predominantly in at most six or seven functional regions, in some cases focusing on as few as two. The relationship between the propagative attractor group ICA component maps and functional regions is not sharp, but is focused enough to render the pdGICA component maps functionally tractable. Temporal correlations between pdGICA component timeseries and net gain/loss functional region timeseries on corresponding windows echoes those traditionally observed between functional region timeseries when aligning each pdGICA component with the functional region with which it has greatest spatial overlap. Schizophrenia strongly disrupts the average correlative relationship between pdGICA components and certain functional regions, some of which tend to be implicated in schizophrenia e.g. the thalamus and the anterior cingulate cortex. Schizophrenia also strongly and pervasively linked to the importance of specific pdGICA components in reconstructing subjects′ observed time-varying propagative density maps. Over half of the 35 pdGICA component make significantly different average contributions to patient propagative density maps than to those of controls, with the functional footprints of impacted pdGICA components spreading over diverse functional domains. Finally, the magnitudes of local directional flows that carry propagation have spatially structured averages and structured, pervasive schizophrenia effects. The framework introduced here follows a new and fundamentally different data-driven approach to the BOLD fMRI signal. We believe that the empirical measurement of local directional flows and wider spatial signal propagation opens a plethora of new avenues through which to investigate healthy and disordered brain function using BOLD fMRI.

## 1. INTRODUCTION

The patterns of activated brain space measured by blood oxygenation-level dependent (BOLD) functional magnetic resonance imaging (fMRI) are typically investigated via pairwise correlations between timeseries corresponding to a fixed collection of functionally-identifiable brain regions or distributed functional networks. In these standard analytical approaches, space – one of the key selling points of fMRI as a source of information about the brain – is at best an implicit consideration, a format into which voxel collections believed to share primary functional roles are mapped. It is ultimately the temporal contributions of these functional nodes on which much subsequent analysis is focused: the spectral properties of individual nodal timeseries, the correlations between pairs of nodal timeseries, correlations computed over full scans (static functional network connectivity (sFNC)) and, increasingly often, also computed on sliding windows through the full scan (dynamic functional network connectivity (dFNC)) [1-32]. Dynamic functional network connectivity can capture large scale patterns of brainwide coactivation that change with time, but this common paradigm yields no information about localized spatial signal flows, nor about the longer-range spatial signal propagation they induce. This persistent blind spot limits our ability to resolve the role of such flows in supporting or impeding the nodal connectivity patterns commonly associated with healthy brain function. The spatiotemporal smoothness of BOLD fMRI creates a data environment in which regional activation growth requires some spatial influx of activation from the region’s immediate neighborhood, and similarly regional activation decline requires drainage of activation from the region into its immediate spatial neighborhood. Observed temporal gains and losses in regional activation are unavoidably intertwined with patterns of signal spatial propagation in the data. While there have been explorations of clearly characterized aerial-view spatial phenomena such as, e.g. traveling waves, in BOLD fMRI [33-46], more granular, data-driven analyses of local directional flows and their larger propagative implications have not been reported in the fMRI literature. The methods we present below are a first step toward addressing a potentially important and long-neglected aspect of the BOLD signal.

## 2. METHODS

### 2.1 Data

We use data from an eyes-closed resting-state functional magnetic resonance imaging (fMRI) study with approximately equal numbers of schizophrenia patients (SZs) and healthy controls (HCs) (n=314, nSZ=151). Scans were preprocessed according to a standard, previously published pipeline [47], with an additional stage of spatial and temporal smoothing to control noise in the estimated spatial derivatives. This extra processing consisted of smoothing preprocessed scans with a 3D Gaussian kernel (*σ* = 3) and 1D temporal moving average with windows of length 3. The gray matter mask for this data contained 60303 voxels: the *x*-dimension (coronal) has length 53; the *y*-dimension (sagittal) has length 63; the *z*-dimension (axial) has length 46. There are 158 sampled timepoints in each scan. All subjects in the study signed informed consent forms.

### 2.2 Spatiotemporal Gradients (STGs)

To diminish confounding of spatiotemporal gradients by subject differences in global signal amplitude, each preprocessed fMRI volume is rescaled by its own mean amplitude. For each voxel *v* = (*x, y, z*) ∈ ℤ^53×63×46^ and each TR t ∈ {1,2, …,158} in an fMRI volume, let ***F***(*v, t*) = ***F***(*x, y, z, t*) denote the amplitude-normalized fMRI signal at (*x, y, z*, ***t***). At every voxel and timepoint, we now consider 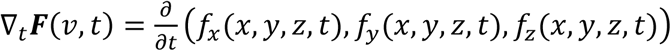 where: *f*_*x*_ = d***F***⁄d*x, f*_*y*_ = d***F***⁄d*y* and *f*_*z*_ = d***F***⁄d*z*. The numerical directional and space-time derivatives are computed via the central difference method, e.g., 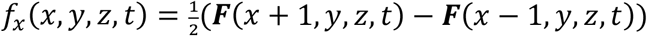 and 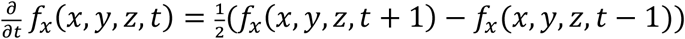. This yields three new volumes 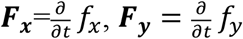 and 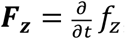, each of the same dimensionality as

### 2.3 Gradient-Driven Propagative Paths and Propagative Path Density Maps

The individual hyperlocal space-time gradients at each 4D point (*v, t*) do not convey much information about propagative patterns in the volume. As a first pass effort to understand the dynamically evolving propagative landscape around each voxel, we investigate gradient driven paths through the brain on intervals of 15TR (30 sec). One path *p*(*v, t*) is initialized at each voxel at the beginning of each 15TR window in the scan (there are 144 such windows, siding one TR at a time). The paths are constructed (**Figure 2** and **Figure 3** (top)) as follows: each voxel has no more than 26 immediate neighbors. There is a unique vector joining the center of the base voxel to the centers of each of its neighbors. The unit normalized space-time gradient *∇*_***t***_***F***(*v, t*) has maximum positive dot product with one of the 26 unit normalized “spokes” connecting ***v*** to its neighbors, i.e. *∇*_***t***_***F***(*v, t*) is maximally directionally aligned toward one neighboring voxel *v′* of *v* at time *t*. So if *p*(*v*, 0) = *v* and *v′* is the neighbor of *v* toward which *∇*_***t***_***F***(*v*, 0) is most aligned, then we set *p*(*v*, 1) = *v′* and subsequently perform the same procedure there using *∇*_***t***_***F***(*v′*, 1) to arrive at *p*(*v*, 2) = *v′′*, etc.

**Figure 1.**
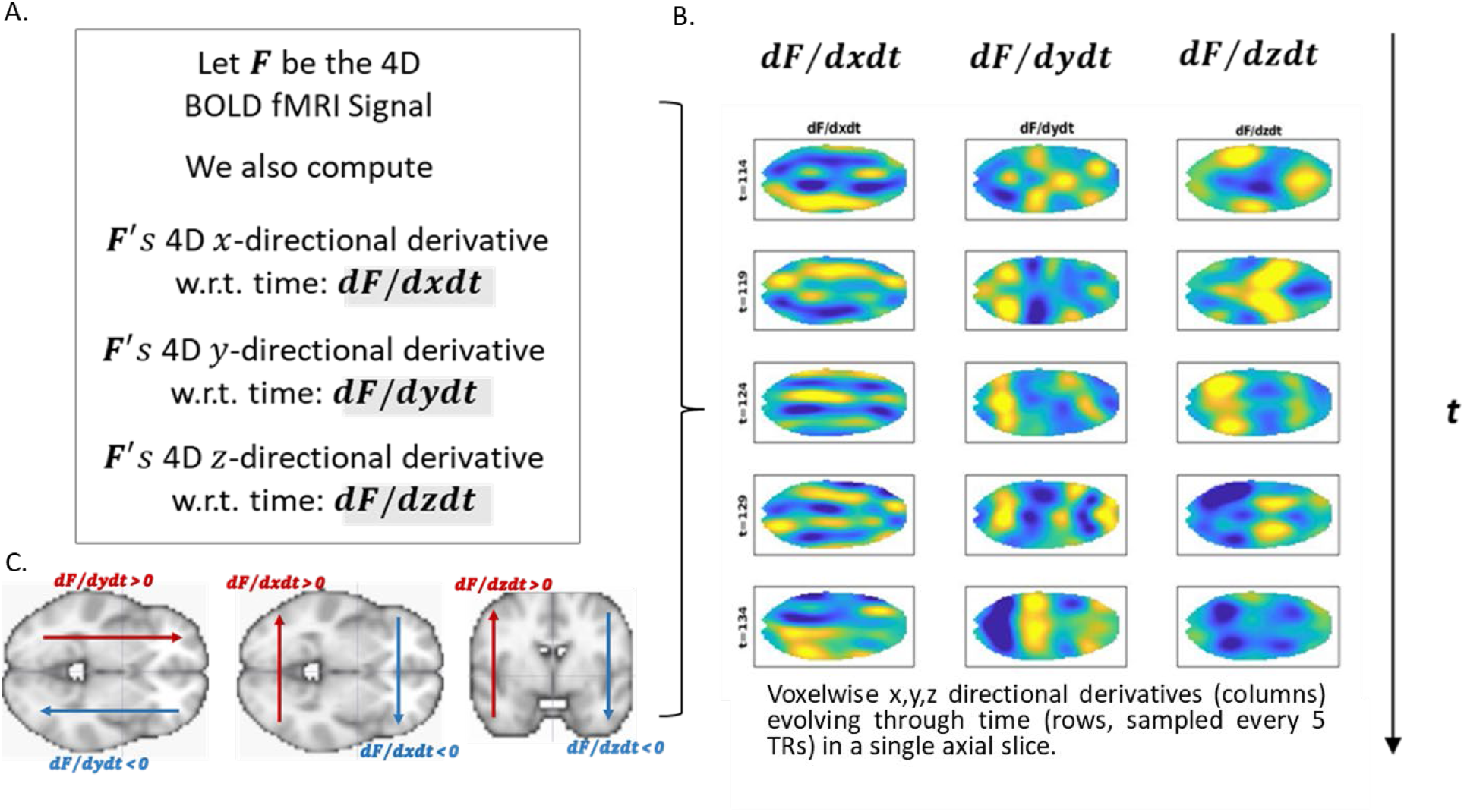
Basic schematic of pipeline. (A, B) with examples of evolving *x*-direction (B, column 1), *y*-direction (B, column 2) and *z*-direction (C, column 3) space-time derivatives; (C) Signed directions along each dimension displayed on axial and coronal brain slices: (C, left) dF/dxdt; (C, middle) dF/dydt (C, middle); (C, right) dF/dzdt, with positive direction along each dimension in red, negative direction in each direction in blue.

**Figure 2.**
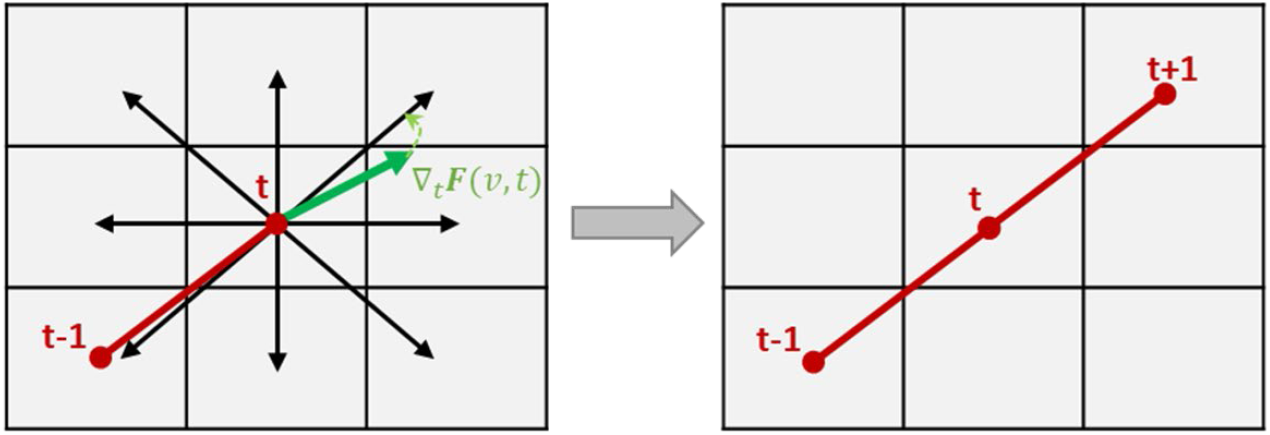
Schematic in planar setting showing how propagative paths are constructed. On the left we see a path *p* entering the center pixel at time ***t*** from the southwest. Vectors joining the midpoint of the center pixel to the midpoints of each of its neighbors are shown as black arrows. The green arrow extending from the midpoint of *p* is the space-time gradient at 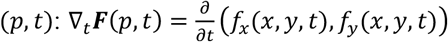 *∇ t* ***F***(*p, t*) is most directionally aligned with the vector connecting *p* to its northeast neighbor *p′*. Thus, the next step on the path, entered at timepoint *t +* 1 is *p*^*′*^, the northeastern neighbor of *p*. Path *p* proceeds in this way through a discrete grid in the plane, driven by the direction of the gradient at each point it visits.

**Figure 3.**
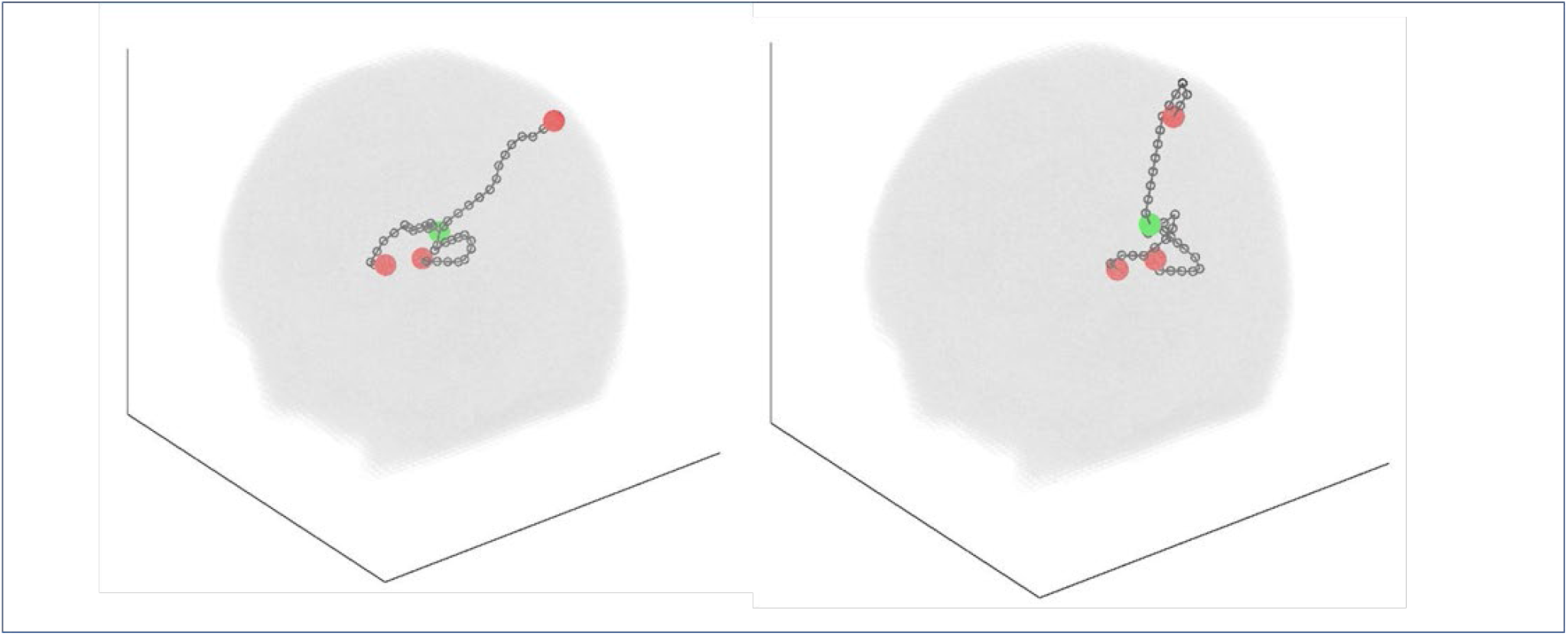
Examples from two subjects (left and right) of three gradient-driven 15-step paths from the green voxel, each initialized at a random timepoint and terminating at one of the red voxels.

We do this for 15 consecutive TRs, starting from each voxel ***v***. This results in 144 paths of 15 TRs 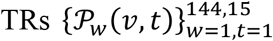 associated with every voxel. For each 15 TR window *w*, running these paths initialized at every voxel creates a distribution of path coordinates on the brain: that is, a *propagative path density (PPD) map*. Each subject has an evolving sequence of 144 PPD brain maps (**Figure 4**) capturing regions and “roadways” in the brain along which spatially propagative energy is concentrated during successive windows.

**Figure 4.**
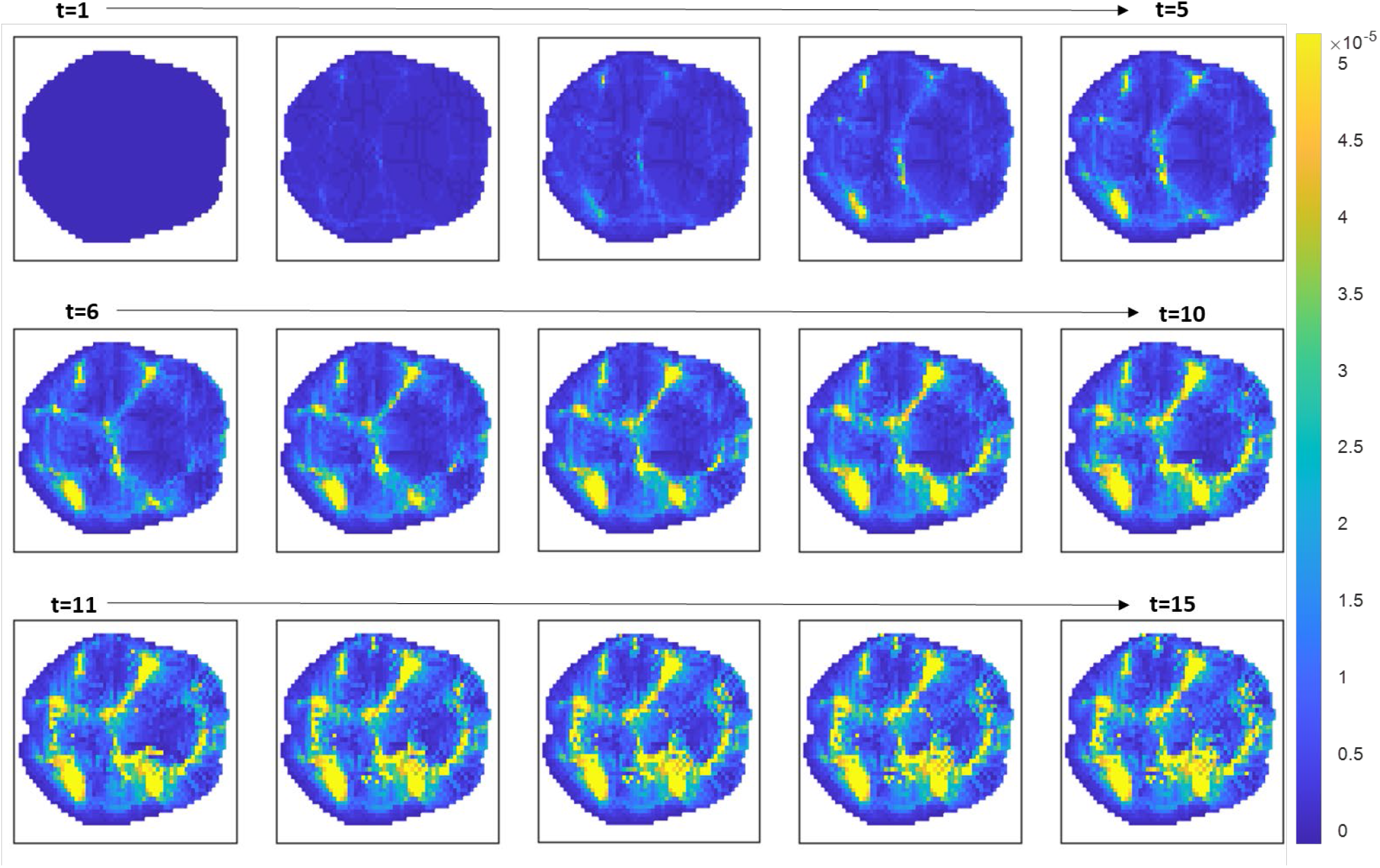
Frame-by-frame evolution of the propagative path density map (on axial slice 18) in one 15TR window. Time goes from left to right, then down. Although the upper bound on densities monotonically increases with time, (as higher and higher levels of concentration in the accumulate in the highest density areas), we are normalizing each frame in this visualization in order to maintain a fixed colorbar. The process is initialized uniformly at t=1, with one abstract “particle” per voxel. Particles then follow paths through the brain, taking one step per TR to a neighbor of their current position determined by the direction of the 3D gradient at the voxel they last landed on at the time they landed there. In this example, the core pattern of concentration is fairly stable by t=10, after which propagative paths evolve primarily along the existing high-density areas, intensifying the pattern. The boxed map in the lower right corner is the PPD associated with the 15TR window being displayed.

### 2.4 Functional Group Independent Component Analysis (fnGICA)

In this work, we utilize output from the functional group independent component analysis (fnGICA)) first reported in [47]. This was an order 100 decomposition, of which 47 components were identified as valid functional networks using criteria elaborated in [47]. The subject-specific timecourses (fnTCs) corresponding to the 47 retained group-level component spatial maps (fnSMs) were detrended, orthogonalized with respect to estimated subject motion parameters and despiked using AFNI’s 3dDespike. The retained fnSMs span 7 functional domains (Subcortical (SC): 5 components; Auditory (AUD): 2 components; Visual (VIS): 11 components; Sensorimotor (SM): 6 components; Cognitive Control (CC): 13 components; Default Mode Network (DMN): 8 components; Cerebellar (CB): 2 components) (**Figure 6**). In much of what follows we will not be using the fnTCs, but rather will employ the corresponding *windowed net gain timecourses* (ΔfnTCs). The ΔfnTCs (**Figure 7**) are simply the summed difference between successive timepoints of the original timeseries on each 15TR window (corresponding to the window used propagative paths detailed above). The original fnTCs have 158 timepoints, and the ΔfnTCs have length 144. This provides a measure of the gain/loss in importance of each functional region to overall signal reconstruction that is concurrent with the windowed measures of propagative path density.

### 2.5 Functionally-Localized Propagative Path Density Maps

Propagative path density maps are computed for each voxel separately in every 15TR window. To assess functionally localized propagative density, we compute the average PPD over the top 5% of voxels in each network on each window, yielding time-varying summaries of the propagative inflow to functional networks (dynNwkPPD). The static version of this measure, summarizing the time-averaged propagative density in each functional network map (nwkPPD) is also investigated here. Note that network thresholding induces disjoint masks by assigning any voxel that is in the top 5% of voxels in more than one network map to the network in which it is maximal.

### 2.6 Propagative Path Density Map Group Independent Component Analysis (pdGICA)

Group independent component analysis was also performed on the time-varying propagative path density brain maps (pdGICA). In contrast to preprocessed fMRI scans, PPD maps are sparse and weblike, representing the cumulative spatial repercussions of local spatially directional signal propagation. The independent components identified by applying GICA to time-varying PPD maps present distributed areas of the brain that tend to concurrently attract multiple propagative paths. In further contrast to fnGICA, the highest-intensity voxels in pdGICA spatial maps (pdSMs) do not necessarily represent tightly coherent sources of the measure of interest, but rather areas onto which propagative paths of unspecified origin tend to converge over 15TR windows. Visual inspection of components produced by pdGICA models of order between 15 and 65 suggests that model orders between 35 and 50 tend to estimate consistent, structured interpretable brain patterns with well-separated high-intensity regions. To keep the analysis relatively compact, we chose to proceed with a model order 35 pdGICA decomposition. It is again worth emphasizing that pdGICA will map the brain differently than fnGICA on the same set of scans. A rough correspondence between pdGICA and fnGICA component maps can be established by aligning each pdGICA component with the functional network with which it shares the largest percentage of high magnitude (top 5%) voxels (see **Figure 16**).

### 2.7 Spatio-Functional Connectivity

This type of analysis has the potential to shed light on how focused patterns of spatial signal propagation relate to observed variations in functional network activation. As a starting point for investigating these prospective relationships, we construct *spatio-functional connectivity* matrices by computing the Pearson correlation between the propagative density map GICA component timecourses (pdTCs) and the windowed net gain timecourses (ΔfnTCs) for each functional network (**Figure 5**, C and D) and (**Figure 7**).

**Figure 5.**
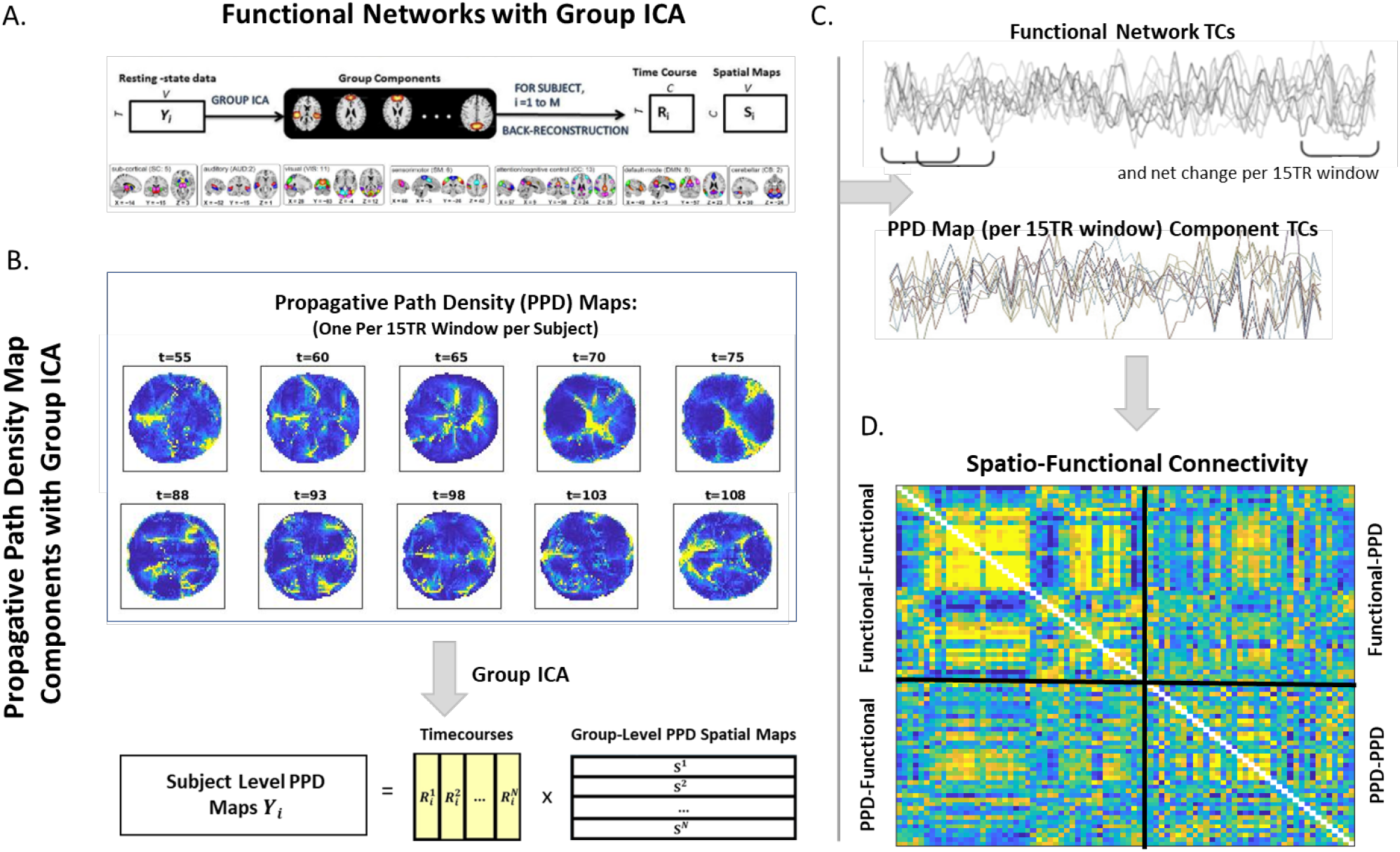
(A) Group ICA (fnGICA) on fMRI scans, producing maximally spatially independent group-level spatial maps (fnSMs) of functional activation and subject-specific timecourses (fnTCs); (C, top); (B) Group ICA (pdGICA) on propagative path density maps producing group-level spatial maps (pdSMs) representing areas that tend to contemporaneously exhibit high propagative density, along with subject-specific timecourses (pdTCs) (C, bottom); (D) Spatio-functional connectivity computed as pairwise correlations between the *net change per 15TR window* in the fnTCs (ΔfnTCs) and pdTCs.

**Figure 6.**
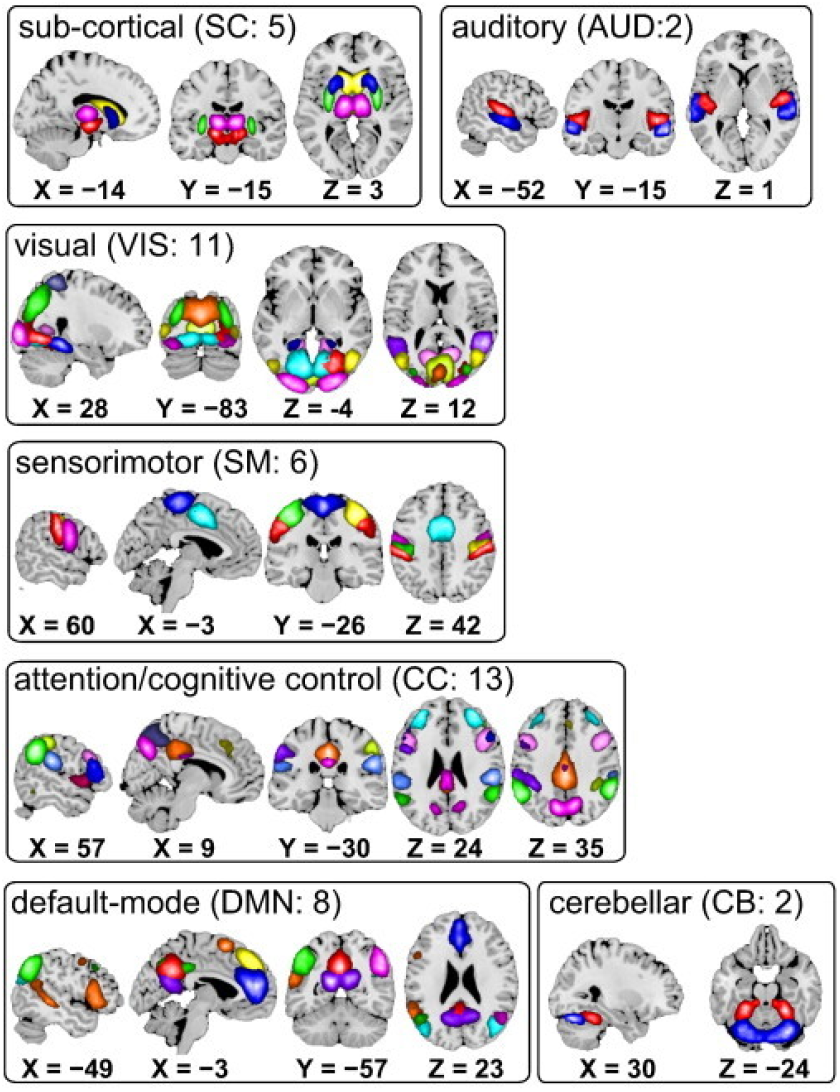
Composite maps of the 47 functional networks, sorted into seven functional domains. Each color in the composite maps corresponds to a different network [47].

**Figure 7.**
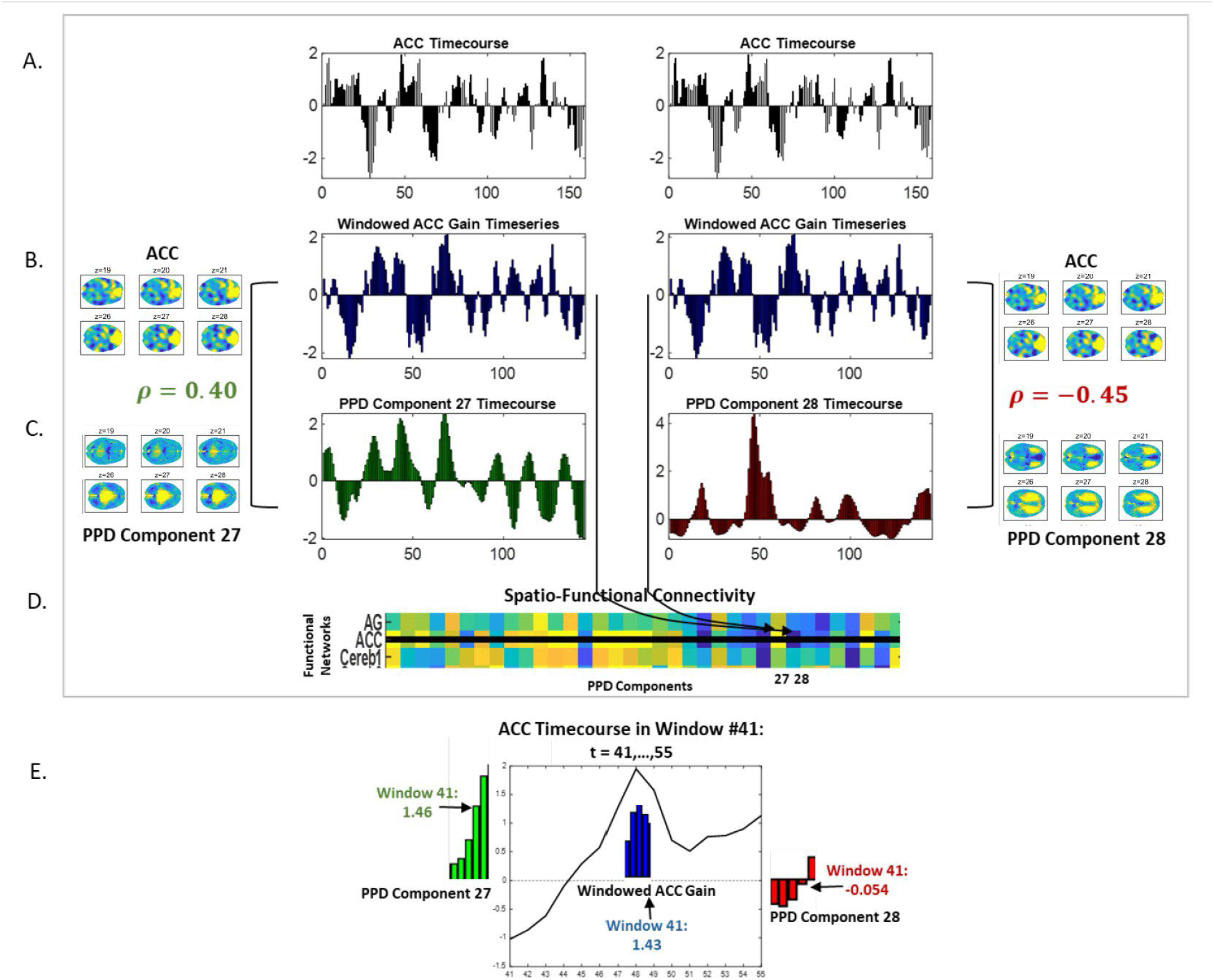
(A) The anterior cingulate cortex (ACC) functional network timecourse for a sample subject (both middle left and middle right); (B, leftmost and rightmost) ACC spatial map, select axial slices (z=19,20,21,26,27,28), smoothed; (B, center left and center right); ACC timeseries of net gain/loss on successive 15TR windows; (C, leftmost) pdGICA component 27 spatial map, same axial slices as in the displayed ACC spatial map; (C, center left) Timecourse for PPD component 27, positively correlated with the windowed ACC timecourse displayed immediately above (C, center left); Timecourse for PPD component 27, positively correlated with the windowed ACC timecourse displayed immediately above; (C, center right); Timecourse for PPD component 28, negatively correlated with the windowed ACC timecourse displayed immediately above; (C, rightmost) pdGICA component 27 spatial map, same axial slices as in the displayed ACC spatial map; (B-C, leftmost) is the positive correlation between ACC net gain timeseries and pdGICA component 27 timecourse, *ρ* = 0.40; ; (B-C, rightmost) is the negative correlation between ACC net gain timeseries and pdGICA component 28 timecourse, *ρ* = *−*0.45; (D) Section from the spatio-functional connectivity matrix of our sample subject. Middle row with black gridline shows subject-specific correlations between ACC net gain ΔfnTC and each of the pdTCs, with components 27 (positive correlation in yellow) and 28 (negative correlation in blue) indicated along the x-axis; (E) Focusing in on a single 15TR window, the 41^st^, showing the key values on that window (E, center) the ACC timeseries within window 41 in black with 41^st^ window net gain shown in blue for short interval of windows inclusive of window 41, on which the value is positive (1.43), i.e., there is a net gain in ACC contribution on that window, consistent with the displayed timeseries; (E, left) PPD component 27 timeseries, which is positively correlated with ACC net gain, showing short interval inclusive of timepoint 41 where its value is positive (1.46) as is the ACC net gain on the same window; (E, left) PPD component 28 timeseries, which is negatively correlated with ACC net gain, showing short interval inclusive of timepoint 41 where its value is negative (-0.054), the opposite sign of ACC net gain on the same window.

### 2.8 Display Conventions

In general, displays of the whole brain are shown as successive axial slices from *z* = 1 (top left) to *z* = 46 (bottom right). Areas in which the time and subject-averaged gradient magnitudes are less than the global median are masked out and displayed in grey. Colormaps are always symmetric about zero when both positive and negative values are being displayed. In the case of strictly non-negative values, colormaps go from smallest non-negative magnitude in cold colors to largest positive magnitude in warm colors. When only six axial slices of the brain are displayed to conserve space, the slices are *z* = 19, 20, 21, 26, 27, 28. When twenty-one axial slices are displayed, the slices are {15, 16, …,35}. Remarks about functional regions with which parts of the display pattern intersect come from importing the volume into MRIcroGL [https://www.nitrc.org/projects/mricrogl/] and employing the AAL atlas overlay.

Displays of functional networks (fnGICA components) along the axis of a figure are always in the same order, organized into functional domains as in **Figure 6** and **Figure 8** (left). Displays of propagative path density components (pdGICA components) along the axis of a figure are always organized so each pdGICA component is aligned with the functional network with which its top 5% of voxels has largest intersection with the top 5% of voxels in that network. Thus reorganized, pdGICAs are displayed in the functional domain containing the functional network with it is best-matched (see **Figure 8** (right)).

**Figure 8.**
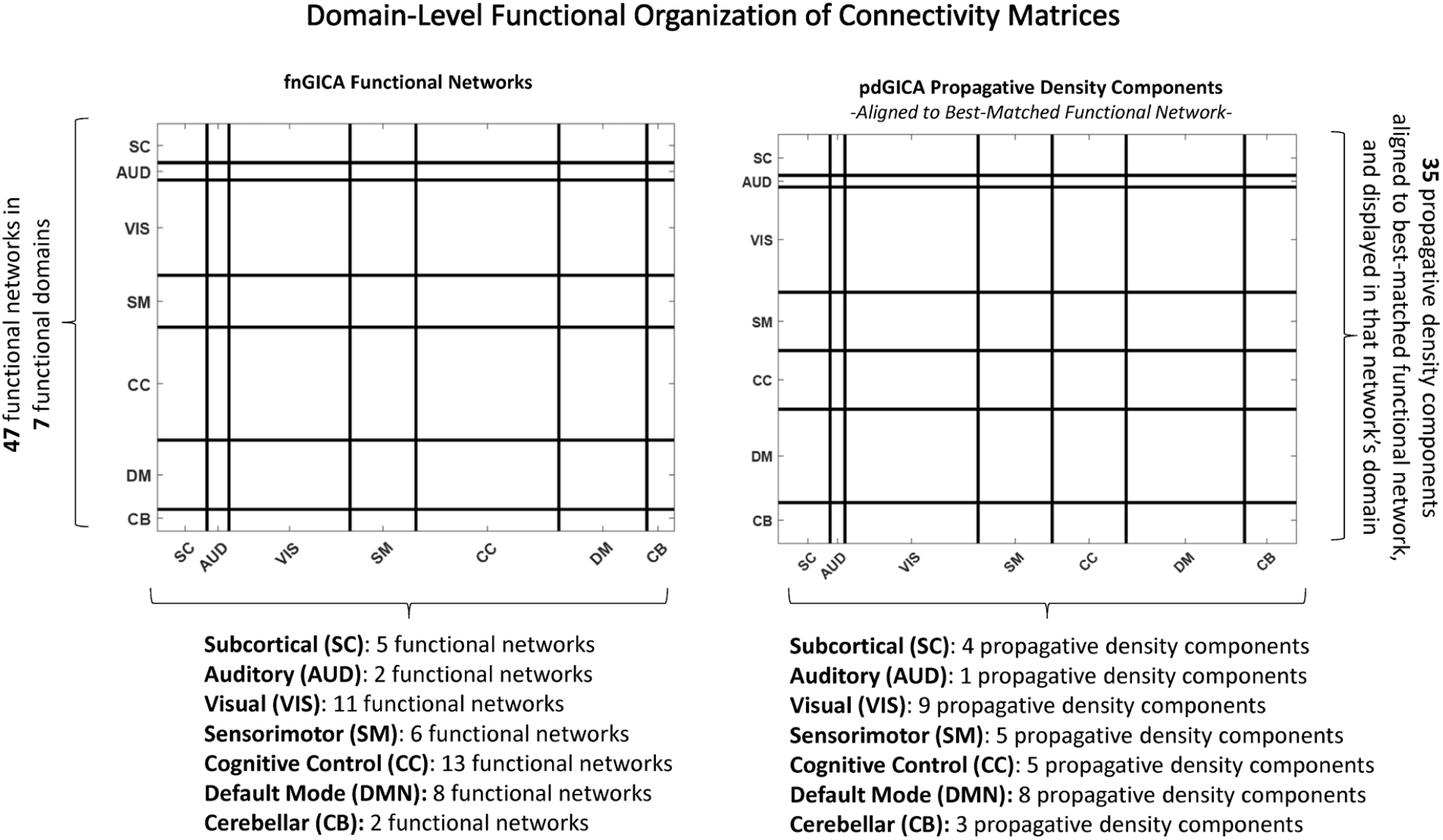
Schematic showing the organization of 47 functional networks (fnGICA components) into 7 functional domains, and presenting the format of domain display along axes (left); also the propagative path density map GICA (pdGICA) analogue of this format (right) in which the 35 pdGICA components are each positioned to align with the functional network with which the top 5% of its voxels have largest intersection with the top 5% of voxels in the network. Then, thus reordered, the pdGICA components are displayed in the domain of the best-matched fnGICA component. The number functional networks in each domain is shown on bottom right, and the number of of pdGICA components whose best-match is a functional network in each of the indicated domains shown on the bottom right.

### 2.9 Statistical Analysis

The SZ effects reported here are obtained through a multiple regression model on SZ diagnosis, gender, age and mean frame displacement. Displayed effects are significant (*p* < 0.05) after correction for multiple comparisons, unless otherwise indicated. Colormaps are symmetric about zero, with cooler (resp. warmer) colors indicating negative (resp. positive) SZ effects.

## 3. RESULTS

We find that local spatial flow strengths averaged over time, both in total magnitude and resolved along individual directions present spatially structured patterns with directionally plausible symmetries (**Figure 9, Figure 10,Figure 11,Figure 12** (left)). The effects of schizophrenia on these measures are also highly structured, clustering in functionally-identifiable areas (**Figure 9, Figure 10, Figure 11,Figure 12** (right)). In general, there are higher magnitude local flow strengths in the occipital areas and propagative path densities are focused more narrowly on a subset of occipital regions, along with temporal lobes, putamen and hippocampus.

**Figure 9.**
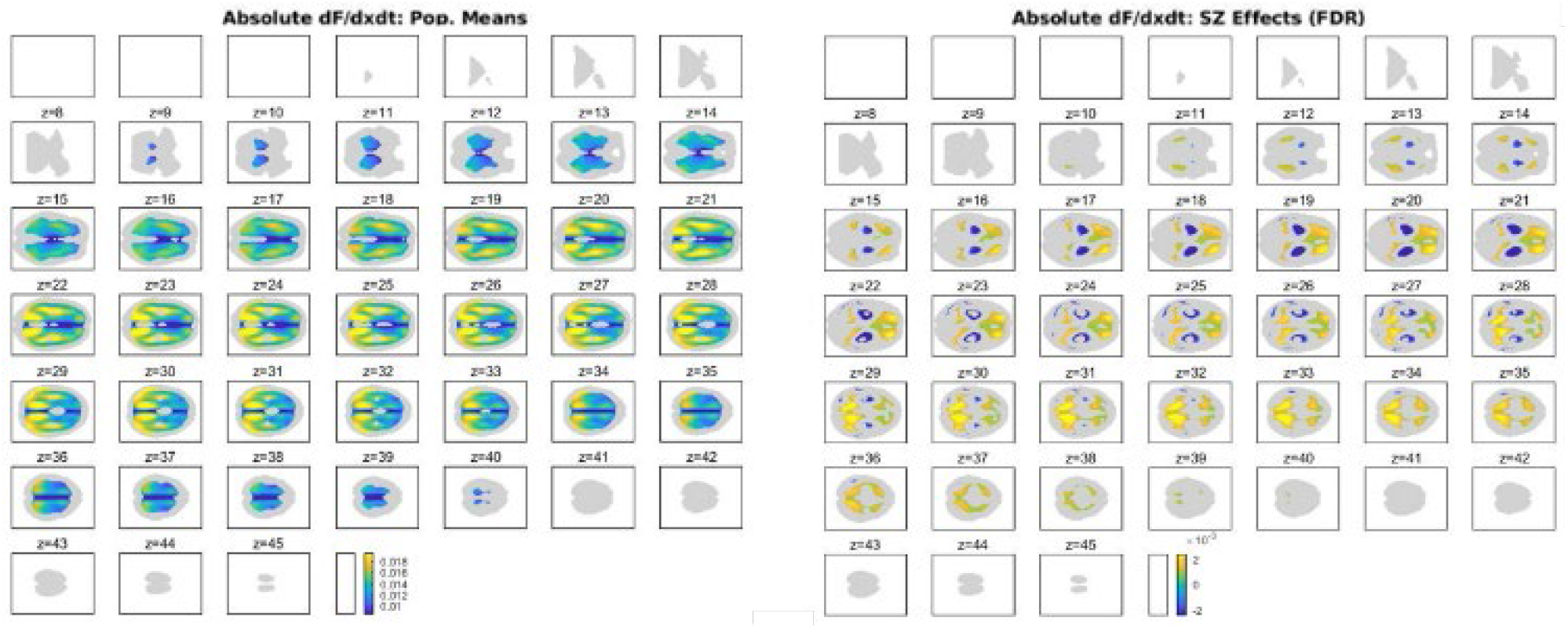
Coronal Direction: Absolute x-Derivative w.r.t time (|*F*_*x*_|). Population means (left) and significant SZ effects (right).

**Figure 10.**
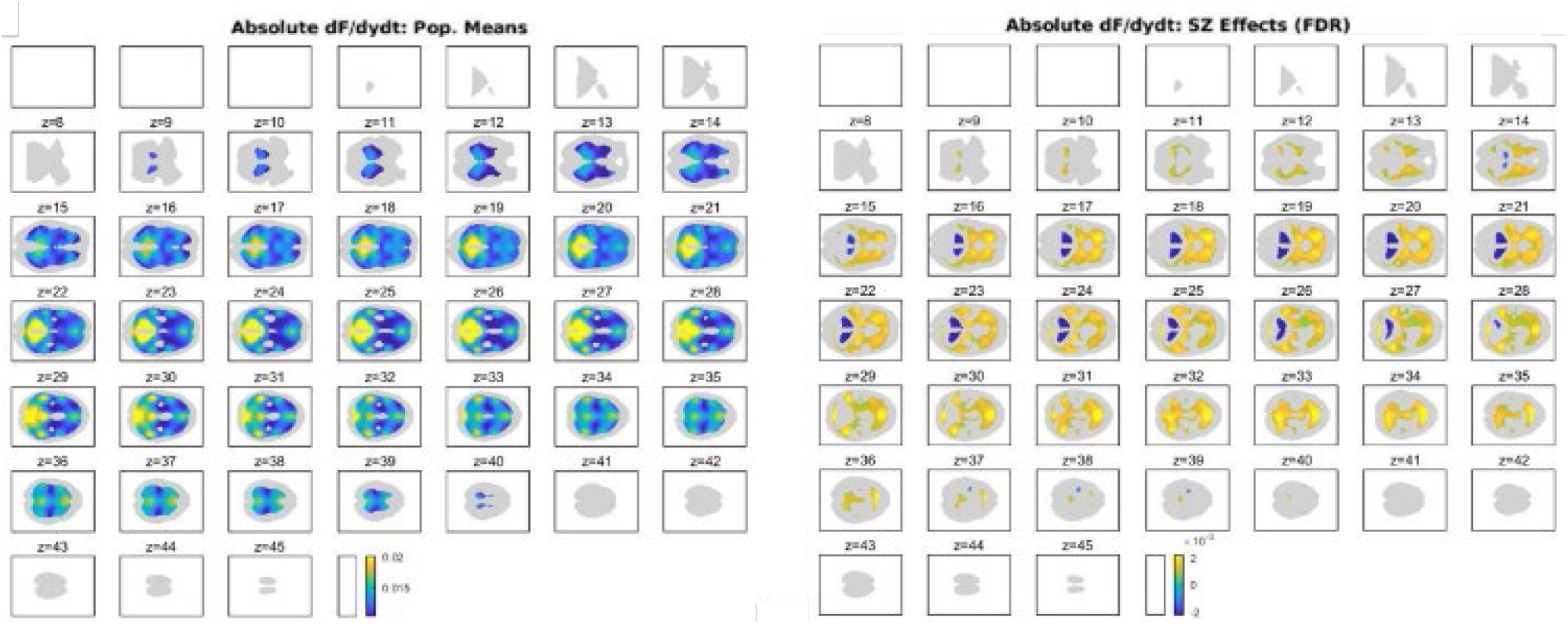
Sagittal Dimension: Absolute y-Derivative w.r.t time (|*F*_*y*_|). Population means (left) and significant SZ effects (right).

**Figure 11.**
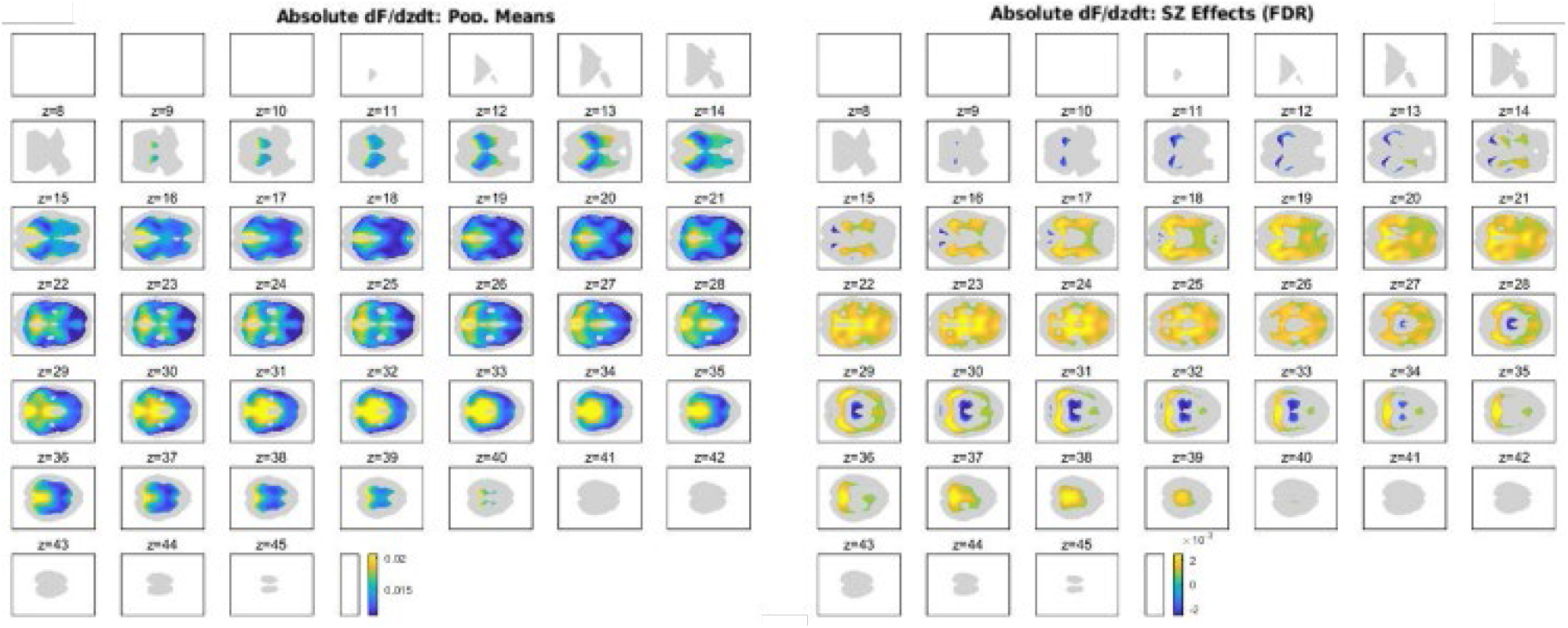
Axial Dimension: Absolute z-Derivative w.r.t time (|*F*_*z*_|). Population means (left) and significant SZ effects (right).

**Figure 12.**
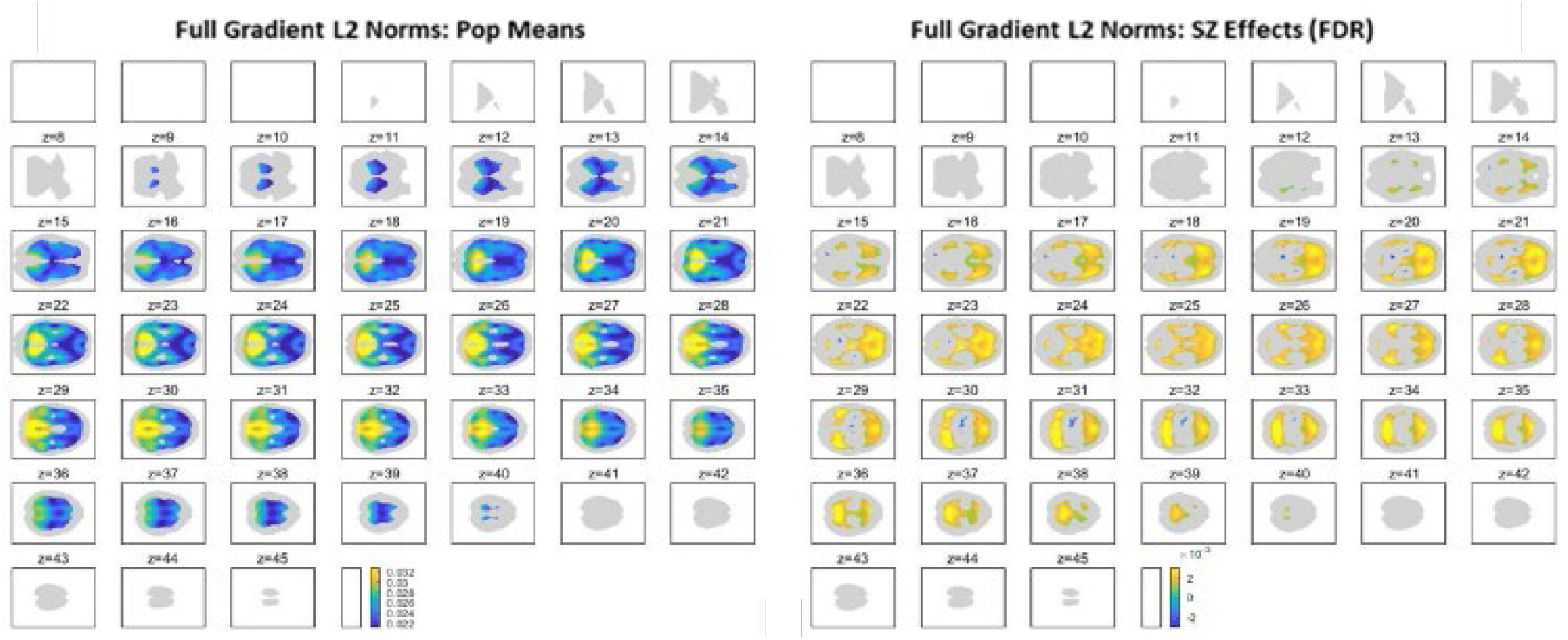
L2 Norms of 3D Gradients (||∇_*t*_*F*(*v, t*)||_2_): Population means (left) and significant SZ effects (right).

### 3.1 Direction-Specific Spatiotemporal Derivative Magnitudes Present Structured Patterns Impacted by SZ

Average voxelwise |***F***_*x*_| and the SZ effects on these magnitudes exhibit bilateral symmetry as one would expect for flows that are directed orthogonally to the corpus callosum (**Figure 9** (left)). |***F***_*x*_| is generally higher in occipital and parietal areas. Schizophrenia tends to further elevate |***F***_*x*_| magnitudes in occipital and parietal regions, and to elevate |***F***_*x*_| in frontal lobes while diminishing hippocampal |***F***_*x*_| (**Figure 9** (right)).

In the case of |***F***_*y*_|, magnitudes exhibit bilateral symmetry and also some symmetries/anti-symmetries along the sagittal plane (**Figure 10** (left)). |***F***_*y*_|, is generally higher in occipital, cerebellar and temporal lobe areas. Schizophrenia significantly elevates sagittal direction local flow strengths in frontal lobes and reduces sagittal direction flow strengths in the cerebellum, with some effects also seen in occipital areas and temporal lobes (**Figure 10** (right)).

Local flows in the *z*-direction (**Figure 11** (left)) are generally stronger in the occipital and temporal lobes than in other parts of the brain, as well as in the thalamus and putamen. Axial slices *z* = 30 through *z* = 34 also present a large proportion of high magnitude local flows along the axial dimension. Schizophrenia pervasively elevates *z*-direction local flow strengths in middle slices *z* = 22 through *z* = 25 (**Figure 11** (right)). Some of this elevation of |***F***_*z*_| intersects with the thalamus, but at higher axial positions, schizophrenia is associated with reduced thalamic |***F***_*z*_|, while SZ significantly increased |***F***_*z*_| in caudate and putamen.

### 3.2 Directionless Summaries of Spatiotemporal Derivative Magnitudes

Unlike the individual direction average magnitudes, the full gradient L2 norm ||∇_*t*_***F***(*v, t*)||_2_ = 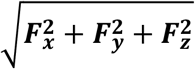 accounts for temporal simultaneity, i.e. it cares that multiple directions have large magnitudes *at the same time*. Aggregated over all voxels, there is a strong anti-correlative trend (*ρ*_*yz*_ = *−*0.29) between *y* and *z* direction local flows. Inspection suggests that the measured anticorrelation arises from high magnitude negative ***F***_*y*_ consistently co-occurring with high magnitude positive ***F***_*z*_ and vice versa. It is worth recalling here from Section 2.2 that positive or negative directional derivatives indicate orientation in space (e.g., ***F***_*y*_ *>* 0 indicates posterior to anterior directionality while ***F***_*y*_ < 0 is flow going in the opposite direction). By contrast *ρ*_*xy*_ = 0.09 and *ρ*_*xz*_ = 0.05. The dominant correlative trend between (opposite sign) magnitudes of ***F***_*y*_ and ***F***_*z*_ is consistent with ||∇_***t***_***F***(***v, t***)||_2_ (**Figure 12** (left)) being highest in regions where both |***F***_*y*_| and |***F***_*z*_| are individually strong. This includes occipital, parietal and cerebellar areas. Schizophrenia effects on ||∇_***t***_***F***(***v, t***)||_2_ are also concentrated where SZ effects are same direction and strong in both |***F***_*y*_| and |***F***_*z*_|. Thus, we see SZ elevating ||∇_***t***_***F***(***v, t***)||_2_ in many occipital, cerebellar, temporal and frontal areas, among others.

### 3.3 Propagative Path Density Maps

Time and subject-averaged voxel-level STG magnitudes cannot reveal information about how flows through each voxel are propagating with time, or how flows through all voxels might be aggregating over the brain through time: e.g., strong magnitudes in neighboring voxels can suggest strong spatial signal propagation in that region or represent flows that double back on themselves repeatedly, without sufficient time or space alignment to progress out of the initial neighborhood. And regions with high average STG magnitudes may not always be part of many different connected propagative pathways such that they play a large role in overall signal propagation during certain windows of time. Propagative density maps, computed on 30 second windows, recover some of this information (see Section 2.3) and for preliminary analysis we present the time and subject averaged means below (**Figure 13** (left)), along with SZ effects (**Figure 13** (right)).

**Figure 13.**
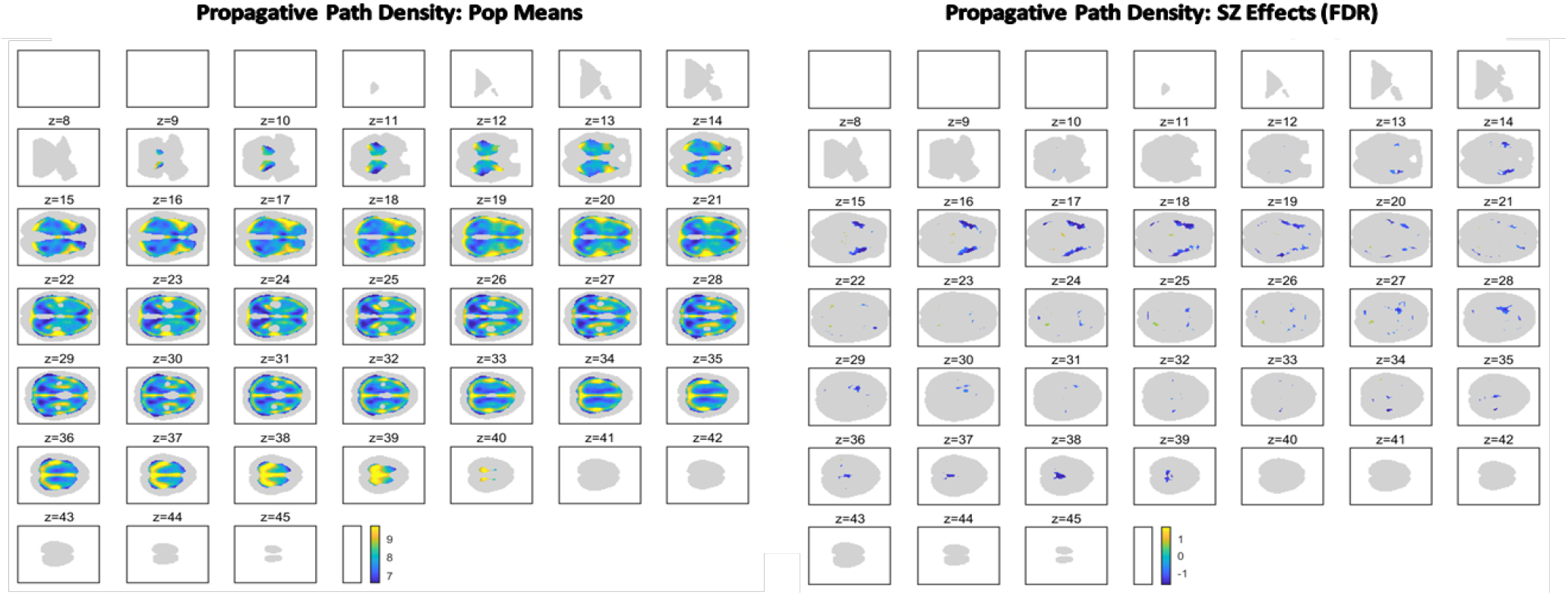
Propagative Path Density Maps. Population means (left) and significant SZ effects (right).

In contrast to the patterns of localized voxel-level flow strengths (directional and directionless), we find that mid-occipital lobes, mid-temporal lobes and cerebellum play a relatively negligible role in carrying longer timeframe spatial signal propagation. This suggests that the high magnitude STGs in those areas are, in the aggregate, repelling: moving signal energy that originates those regions out relatively quickly and not attracting propagation from other parts of the brain. Schizophrenia affects this measure less pervasively than the local STG magnitudes, significantly lowering PPD in only a few areas, including the angular and fusiform gyri and small sections of cerebellum, the frontal lobe, amygdala and insula.

### 3.6 Functionally-Localized Propagative Path Density Maps

By averaging voxel-level propagative path density within functional network masks we can examine the functional differentiation of propagative flow into brain areas. Network masks consist of the top 5% of voxels in each functional network spatial map. In cases where a voxel meets this criterion in more than one network, it is assigned to the network in which its z-scored value is largest. Time-varying, or “dynamic” network-masked PPDs (dynNwkPPDs) are computed as the average within-mask PPD on each 15TR window. From this, we also obtain a time-averaged summary of PPD within network masks (nwkPPD).

One question is whether certain functional areas have a greater tendency to collect or attract BOLD signal propagation, while others either actively repel spatial flows or act as transient channels for propagation being drawn toward other parts of the brain. And if such patterns exist, do they differ in systematically between healthy populations and subjects with severe mental illnesses such as schizophrenia. We uncover evidence that spatial signal flows, generally have a greater tendency to accumulate in interior and posterior brain regions (see **Figure 14** (Top Row)), concentrating especially in the posterior cingulate cortex, cuneus, precuneus, medial occipital gyrus, lingual gyrus, calcarine, superior temporal gyrus and posterior insula. Notably, schizophrenia elevates propagative concentration in a set of frontal areas, mostly related to the inferior frontal gyrus, and two key default mode networks (posterior cingulate cortex and dorsomedial prefrontal cortex) (see **Figure 14** (Bottom Row)). And while propagative path density in PCC is high across the whole population, it is further elevated in schizophrenia. The IFG and dMPFC also have significantly elevated nwkPPD in schizophrenia patients, but in contrast to the case of PCC, these networks have extremely low levels nwkPPD on a population-wide basis.

**Figure 14.**
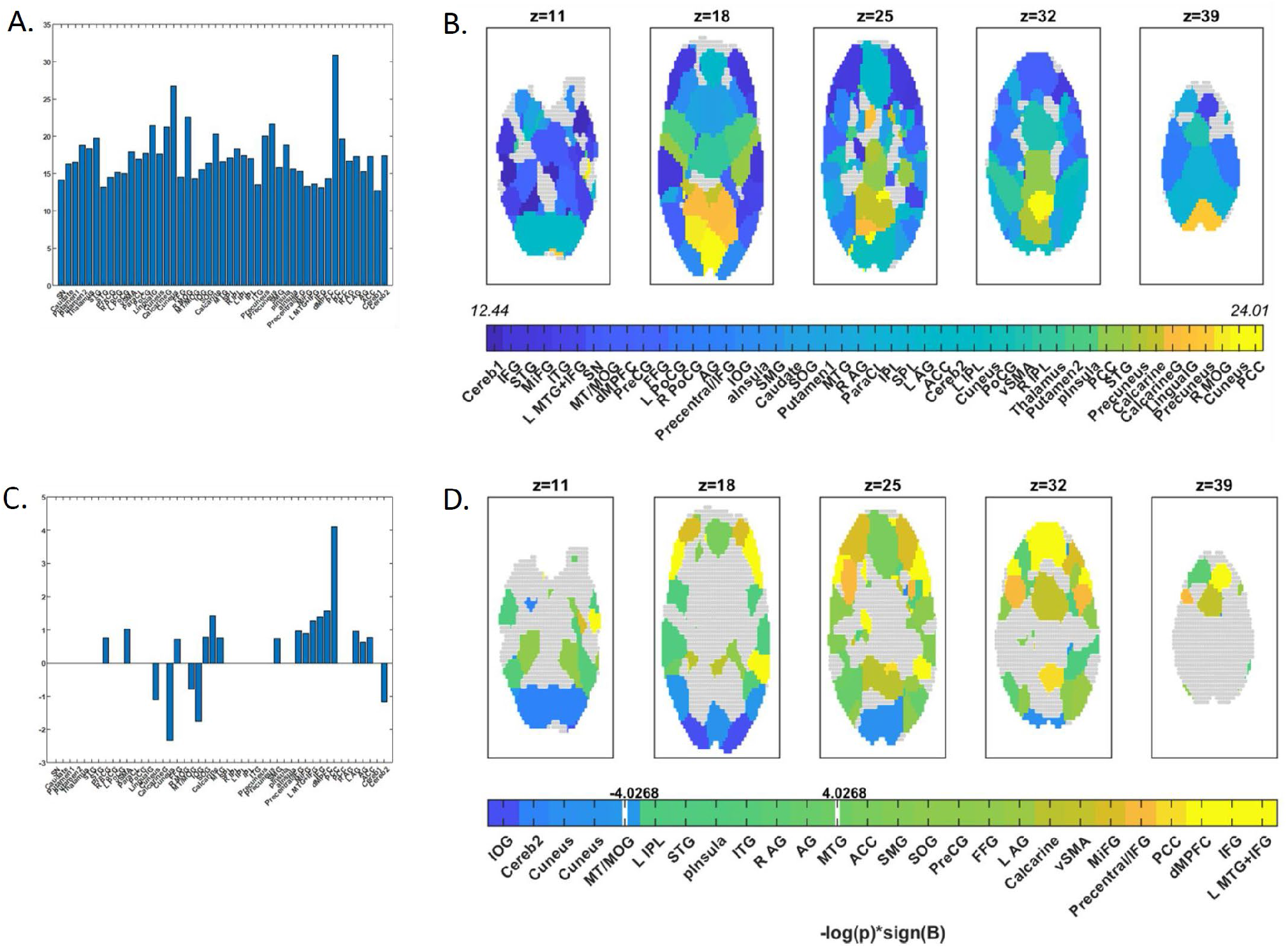
Time-averaged propagative path density (nwkPPD) within functional network masks; (A) Population means displayed for each network in bar graph form with networks arranged in the standard format along the x-axis (see Section 2.8); (B) Population means displayed spatially in select ascending axial slices (z=11, 18, 25, 32, 39), colorbar includes only colors that map to network means and labeled with corresponding network names in order of lowest to highest; (C) Significant (p<0.05 (FDR)) SZ regression effects displayed for each network in which the effect is significant after correction for multiple comparisons in bar graph form with networks arranged in the standard format along the x-axis); (D) Significant SZ regression effects expressed as *−* log(*p*) sign(*β*) in select ascending axial slices (z=11, 18, 25, 32, 39), colorbar includes only colors that map to significant (p<0.05 (uncorrected)) regression effects and is labeled with corresponding network names in order of lowest to highest; signed bounds for FDR-corrected significance indicated with white vertical bars and numerical labels.

In principle, one might expect propagative BOLD influx to functional brain areas to correlate positively with the activation growth (the ΔfnTCs) in those areas. This expected pattern holds across all networks (**Figure 15** (C)), with thalamus and calcarine gyrus presenting the strongest correlations between nwkPPD and net-grain, and inferior occipital gyrus and cerebellum presenting the lowest. The strong correlation between nwkPPD and net-gain in the thalamus is significantly diminished in schizophrenia patients (**Figure 15** (D)) while the correlations between nwkPPD and net-gain in several frontal and default mode networks is elevated in patients. Across networks we see that nwkPPD in many non-visual regions is, on average, anti-correlated with net-gain in visual networks (**Figure 15** (A)) and that generally the correlation between nwkPPD and net-gain for network-pairs in the same functional domain tends to be lower than correlations between net-gain timeseries themselves within functional domains (e.g., compare **Figure 15** (D)) with (**Figure 18** (left))

**Figure 15.**
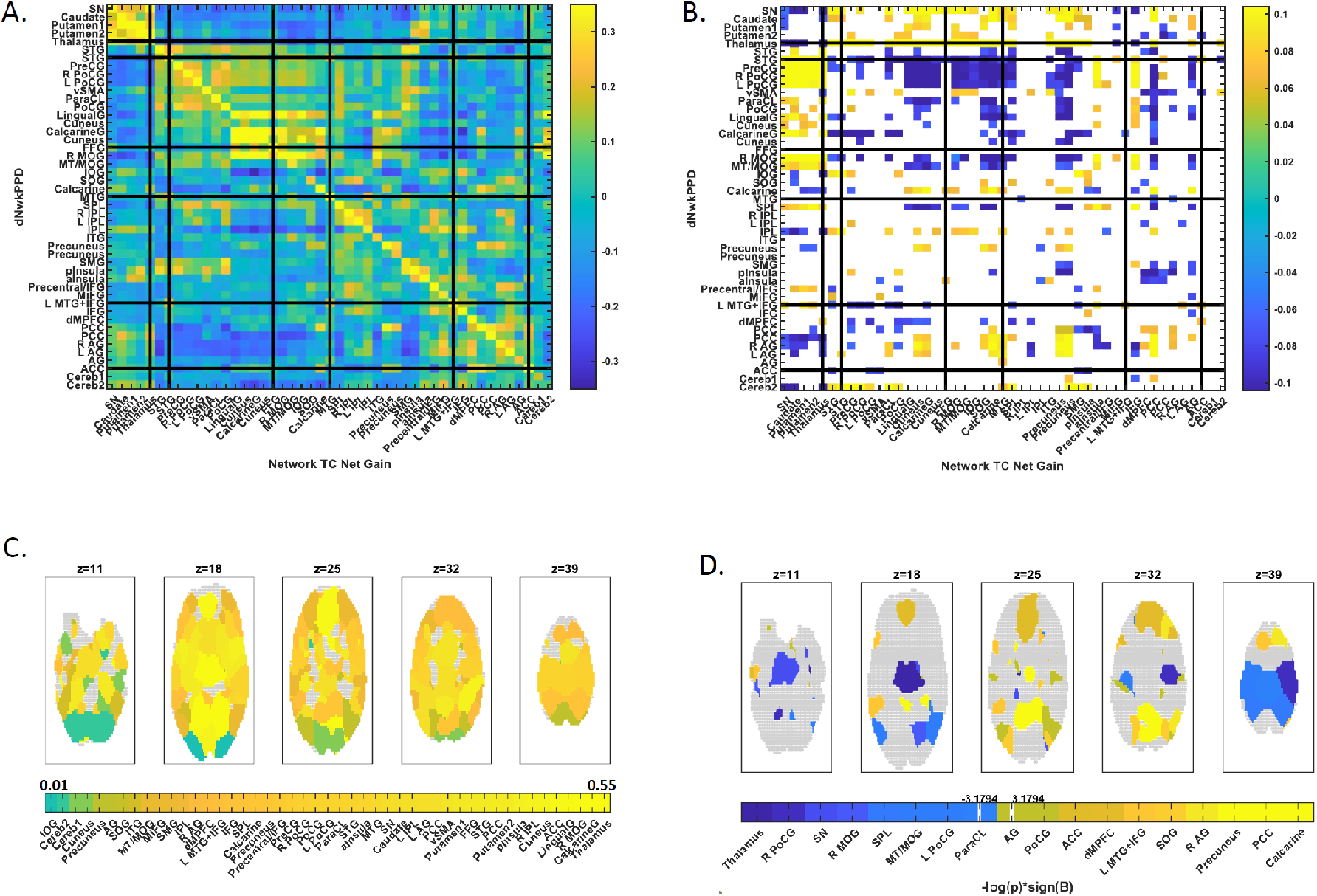
(Top Row) Correlations for all network-pairs, between time-varying functionally-localized network-masked PPD (dynNwkPPD) and windowed network-level net-gain TCs (ΔfnTCs); (A) Population means; (B) Significant effects of SZ (p<0.05 (FDR)); (Bottom Row) Correlations for all individual networks between time-varying functionally-localized network-masked PPD and ΔfnTCs. This corresponds to the diagonals of the heat maps in the top row, but isolates the degree of correspondence between dynNwkPPD and ΔfnTCs for each individual network. One expects these correlations to be positive, in general, as the spatial inflow of BOLD signal to a region would support growth in regional activation (C); Population means displayed spatially in select ascending axial slices (z=11, 18, 25, 32, 39), colorbar includes only colors that map to network means and are labeled with corresponding network names in order of lowest to highest value; (D) Significant SZ regression effects expressed as *−* log(*p*) *sign*(*β*) in select ascending axial slices (z=11, 18, 25, 32, 39), colorbar includes only colors that map to significant (p<0.05 (uncorrected)) regression effects and is labeled with corresponding network names in order of lowest to highest; signed bounds for FDR-corrected significance indicated with white vertical bars and numerical labels.

### 3.7 Propagative Path Density Map Group ICA (pdGICA)

The pdGICA spatial maps are highly structured and although they rarely localize within specific functional networks, they are often focused primarily in one or two functional domains (see Methods, Subsection 2.6 and **Figure 16**). There are pervasively significant differences between SZ patients and controls in the contributions these components make to reconstructing subject PPD maps (**Figure 17** (B)). Of the six pdGICA components that make significantly larger contributions to SZ patient PPD maps, half (components 4, 5 and 13) are spatially focused on DMN networks. The six pdGICA components with the strongest negative SZ effects (components 15, 19, 21, 27, 28, 30) include two (components 27 and 28) that are spatially focused on DMN networks, with the other four presenting spatial foci spanning subcortical, auditory, visual and cerebellar regions. In addition to the functional domains within which pdGICA components spatially focus, the components present other distinguishing characteristics, e.g. some are lateralized (component 5) while others are bilaterally symmetric (components 30 and 31). Bilateral components are more common, suggesting the propagative concentration on both sides of the corpus collosum may frequently be harmonized by ambient cognitive, attentive or behavioral states of the subject since hyperlocalized directional flows around, e.g. two spatially distal bilaterally symmetric voxels, would not be influencing each other directly.

**Figure 16.**
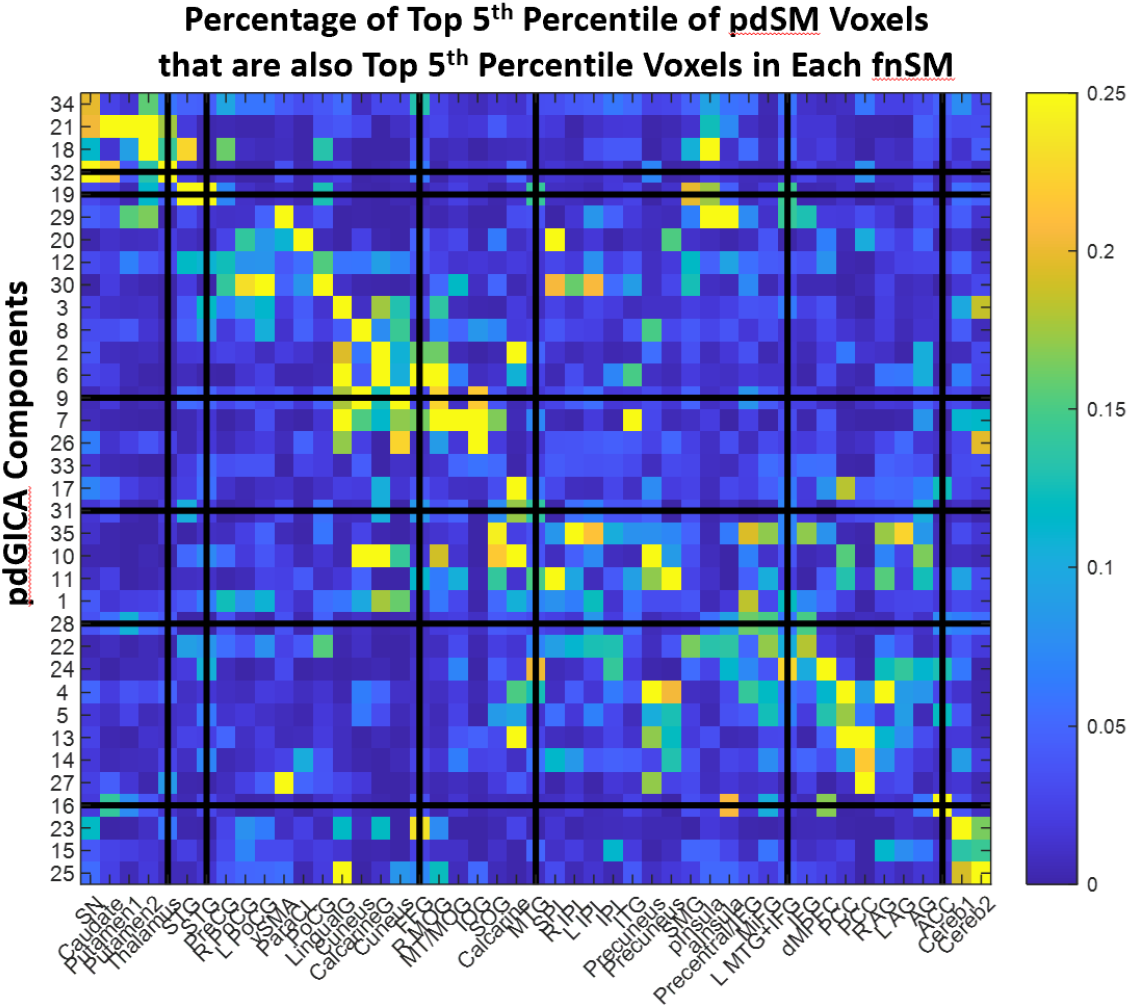
Rows display the percentage of the upper 5^th^ percentile of voxels in each pdSM that intersect with the set of upper 5^th^ percentile voxels in each fnSM. The pdSMs are arranged along the y-axis according to the fnSM with which they have maximal overlap. Some pdSMs are concentrated on one domain or a pair of domains, e.g. pdSMs 21 and 32 are focused on subcortical networks; pdSM 30 spans visual and cognitive control domains; pdSMs 2, 3, 6, 7, 8 and 9 are focused in a set of visual networks, with some including networks in the sensorimotor domain. Other pdSMs are highly focused on specific networks: e.g., pdSM 14 is concentrated on the posterior cingulate cortex (PCC), or on functionally distributed network pairs/triads: e.g., pdSM 16 distributes over anterior insula (aInsula in Cognitive Control), PCC and anterior cingulate cortex (ACC) in default mode network (DMN); while others are more evenly spread over fnSMs: e.g. pdSMs 6, 12, 22 and 31. Mapping the brain into spatially independent components based on the concurrent propagative importance of voxels dimensionally reduces the fMRI signal in novel ways, producing flow-based component maps that span multiple functional regions.

**Figure 17.**
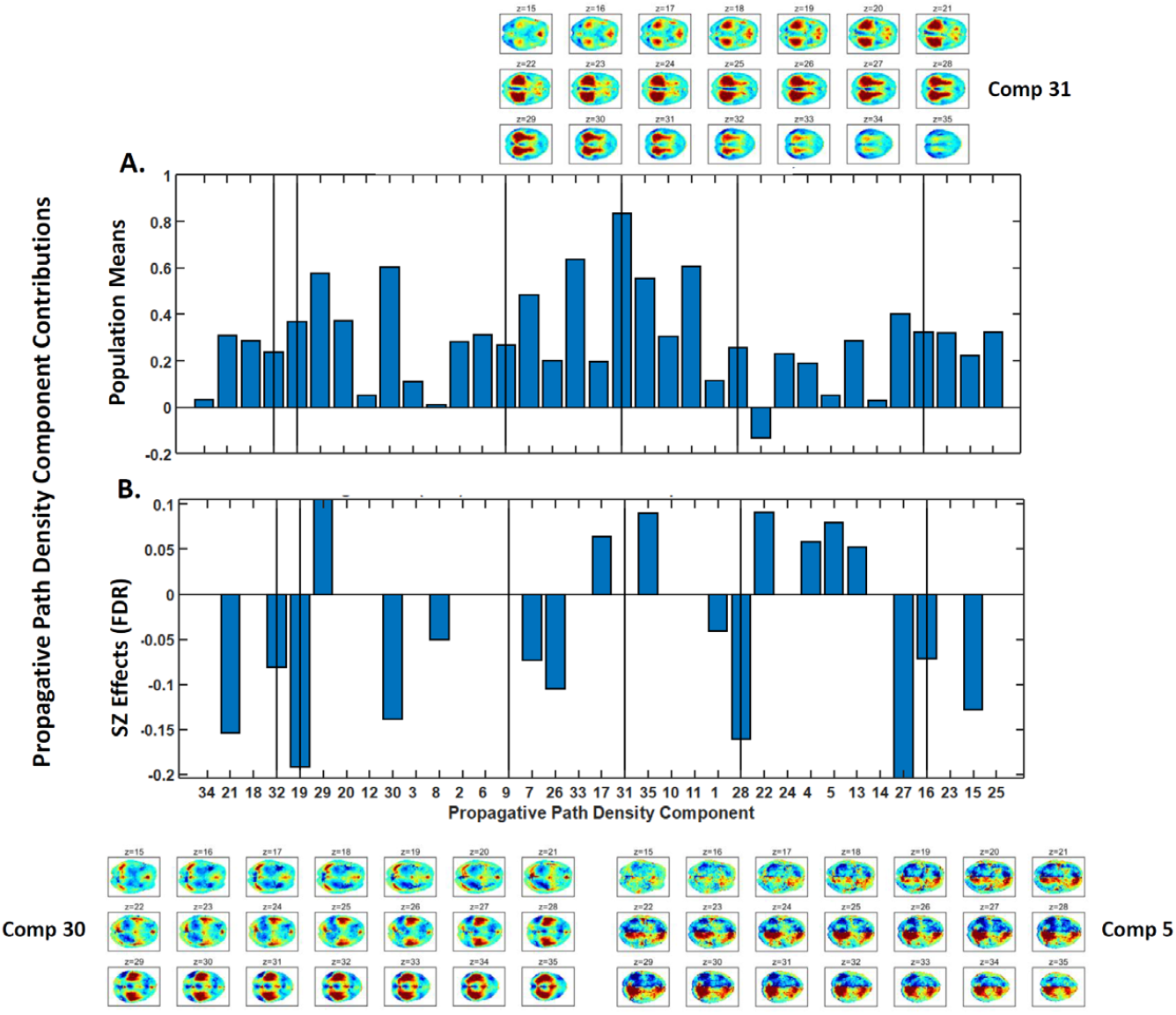
(A) Mean contribution of each pdGICA component. Component 31, which distributes relatively evenly over functional regions (see **Figure 16**), has the highest average contribution; axial slices 15 through 35 of Component 31 are displayed above the bar plot of means; (B) SZ effects on the mean contribution of pdGICA components. Component 5 -a highly lateralized component, exhibiting substantial overlap with right angular gyrus (see **Figure 16**) - makes significantly larger contributions to patient PPD maps, and Component 30 – overlapping primarily with a subset of visual and cognitive control networks - makes significantly smaller contributions to patient PPD maps; axial slices 15 through 35 of Component 5 and Component 30 displayed in the bottom right and bottom left respectively.

### 3.8 Spatio-Functional Connectivity

#### 3.8.1 Functional Network Connectivity

Functional network connectivity (FNC) is a very standard measure computed from resting state fMRI. It is typically assessed as the Pearson correlation between functional network timecourses, and for this study we have adapted the convention slightly by using the windowed net-gain network timecourses, ΔfnTCs, to facilitate comparisons with the spatio-functional connectivity between pdGICA component timecourses and ΔfnTCs reported below. The patterns of connectivity and of significant SZ effects on this connectivity (**Figure 18**) is very similar to that reported using correlations between the fnTCs themselves [4, 28, 47-52] rather than the derived windowed net-gain timeseries

**Figure 18.**
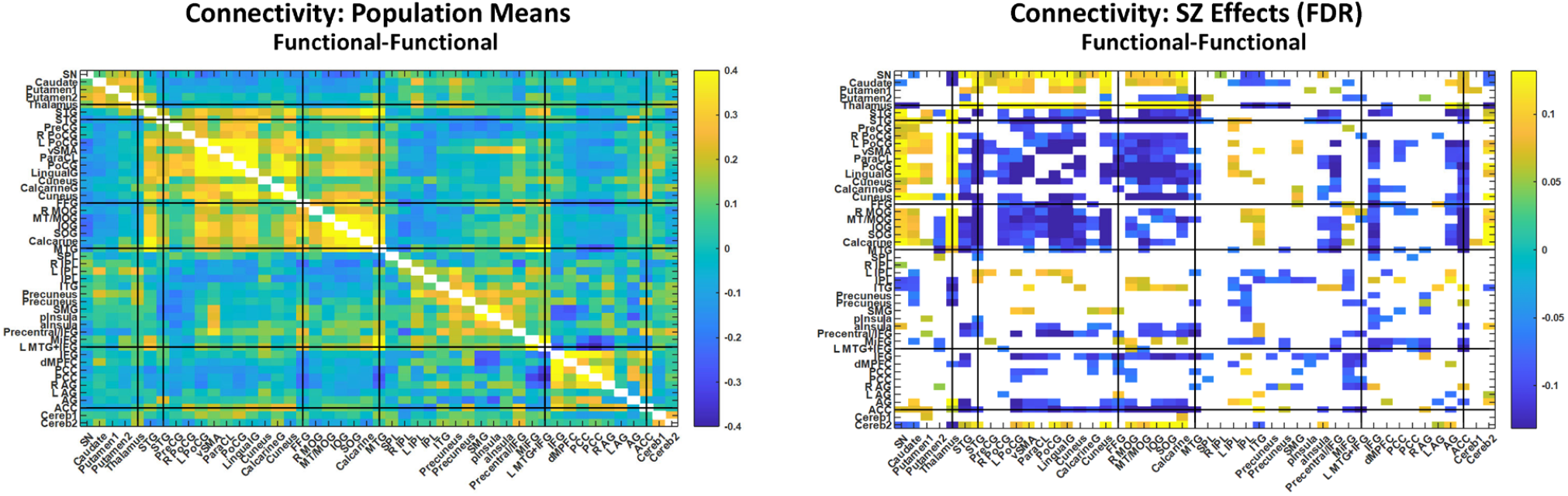
Functional network connectivity computed as the Pearson correlation between ΔfnTCs; (Left) population means; (Right) Significant (*p* < 0.05 (FDR)) SZ effects.

#### 3.8.2 Spatio-Functional Connectivity

One of the main motivations for studying spatial signal propagation in BOLD fMRI data is to explore the role that local spatial flows may have in supporting or impeding healthy functional activation patterns. The *spatio-functional connectivity* between pdGICA and fnGICA components is one way to capture relationships between propagative and functional activity in the brain. Patterns of positive and negative correlations between pdTCs spatially focused in certain functional areas and actual ΔfnTCs (**Figure 19** (Right)) resemble those observed between the ΔfnTCs themselves (**Figure 18** (Left)).

**Figure 19.**
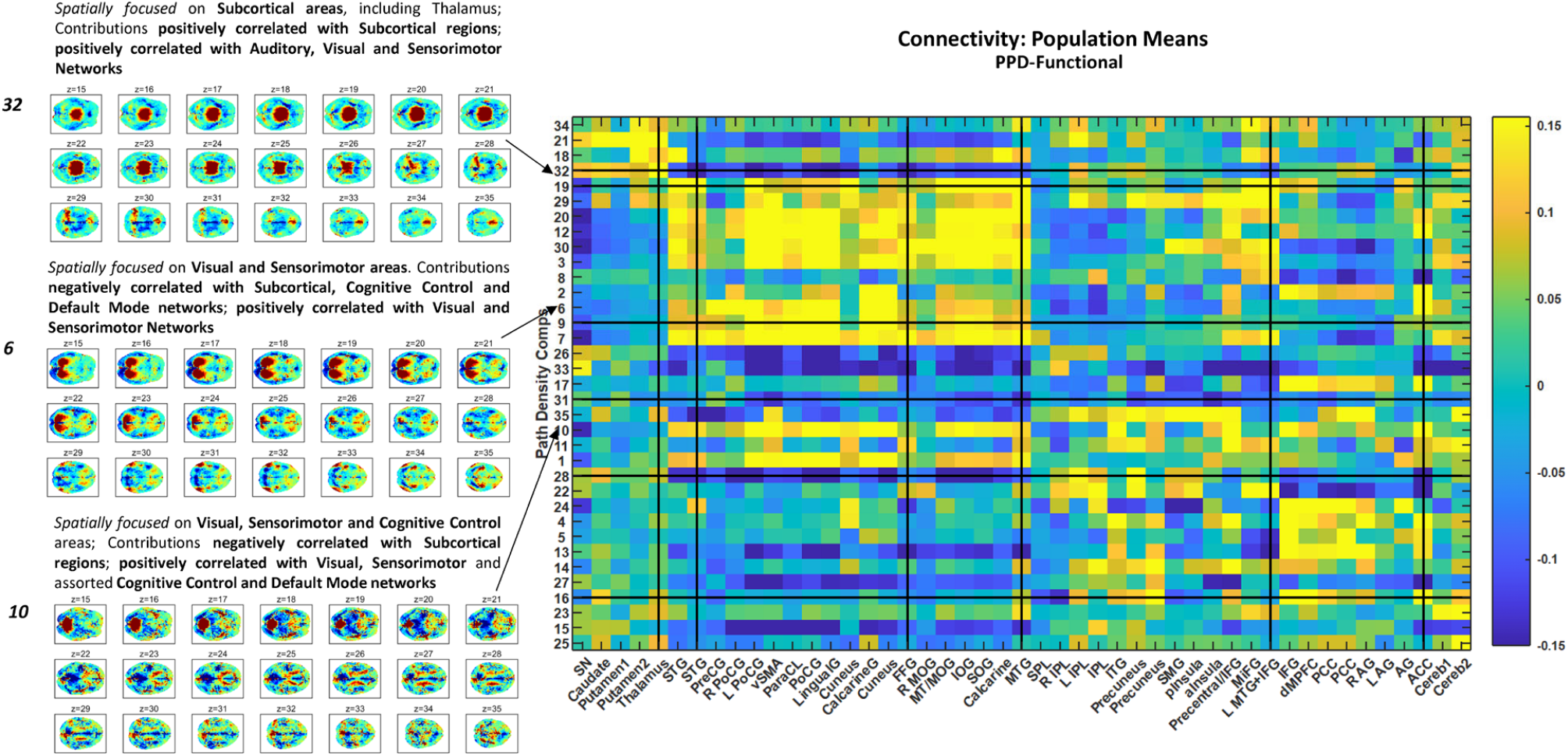
(Rightmost column) Population average spatio-functional connectivity. The 35 pdGICA components are displayed on the y-axis, arranged in order of the fnGICA maps to with which they have largest overlap (as in **Figure 16**) and the corresponding 47 labeled fnGICA components are displayed on the x-axis. The (i,j)^th^ matrix element contains the average correlation between subject-specific timecourses for the pdGICA component in position i on the y-axis, and the net gain timeseries for fnGICA component in position j on the x-axis. (Left Column) Select axial slices (z=15 through z=35) of component maps for three pdGICA components whose temporal correlations with fnGICA component ΔfnTCs are significantly impacted by SZ (see **Figure 20**); pdGICA components 32 is spatially focused in subcortical areas and figure suggests that propagative density in that area is, on average, correlated positively with activation patterns; pdGICA components 6 and 10 are spatially focused in visual and sensorimotor areas and also present evidence of correlation between propagative accumulation and activation in visual and sensorimotor areas.

We see that schizophrenia most significantly impacts the relationships of propagative density distributions with a select group of functional networks (**Figure 20** (Right)), including but not limited to: the substantia negra, the thalamus, the left postcentral gyrus, the medial temporal gyrus, the left MTG and inferior temporal gyrus and the anterior cingulate cortex. Of these, it is the thalamus that exhibits the largest number of significantly relationships with pdGICA components that are significantly disrupted by schizophrenia.

**Figure 20.**
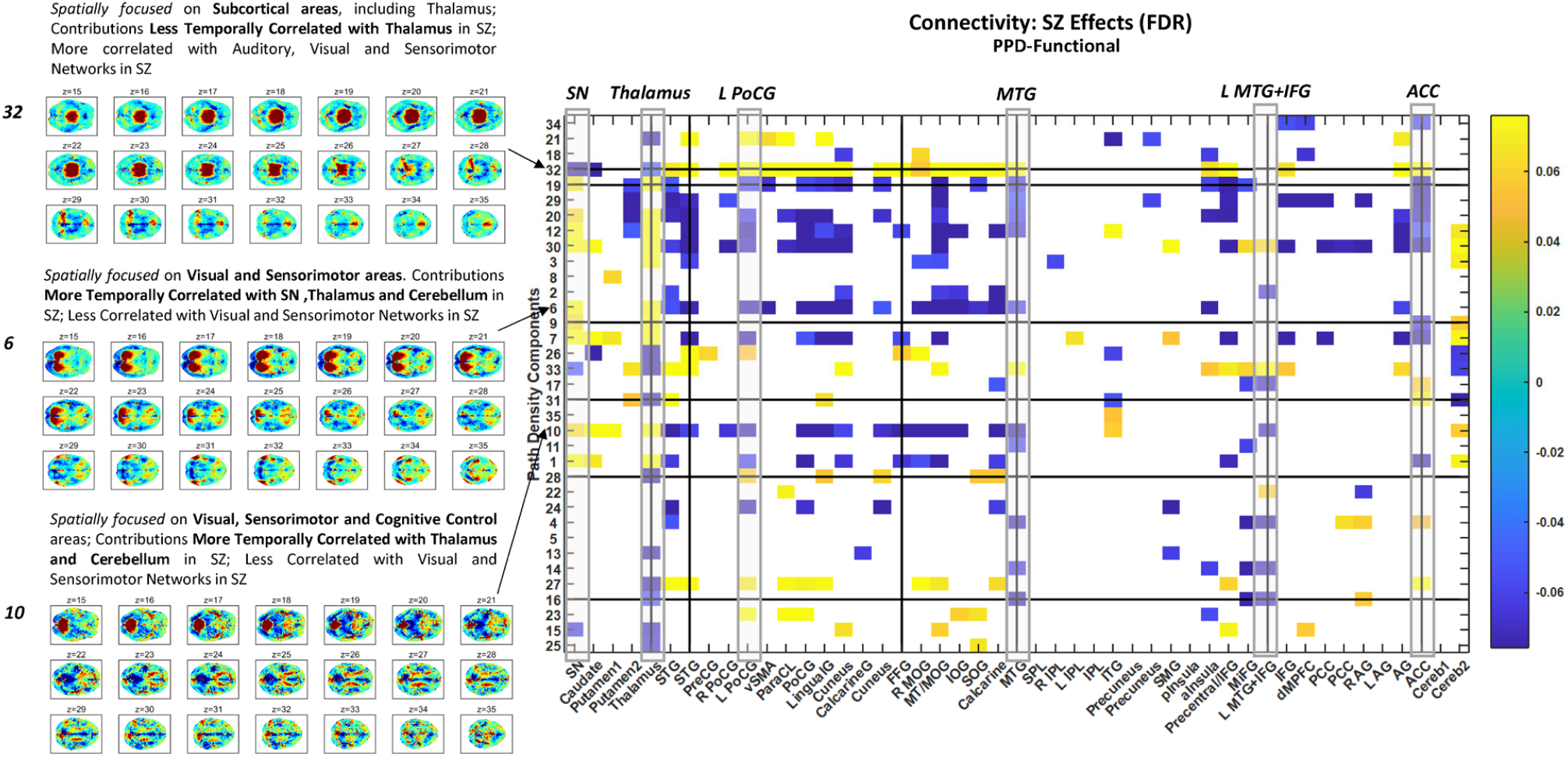
(Rightmost column) SZ effects on spatio-functional connectivity. SZ effects on the correlations between propagative density GICA components and functional GICA components are focused on a subset of functional networks, including (highlighted in grey): the substantia negra, the thalamus, the left posterior central gyrus, the left medial temporal gyrus, the inferior frontal gyrus and the anterior cingulate cortex. SZ effects on the correlations between pdGICA components and functional networks also focus on certain pdGICA components, among which are components 32, 6 and 10, each displayed on the left, exhibited on axial slices 15 through 35.

#### 3.8.1 Spatio-Propagative Connectivity

The correlative relationships between pdGICA component TCs, a type of *spatio-propagative connectivity*, provides information about regions of brain that that tend to concurrently gain (resp. lose) propagative signal concentration vs. those which may be relinquishing concentration as propagation density accumulates in other parts of the brain. One notable feature of the spatio-propagative connectivity in this study is that apart from pdSMs that are spatially focused on visual areas, the connectivity between pdSMs focused within a given functional domain tends to be considerably weaker and more heterogeneous than the corresponding functional network connectivity or spatio-functional connectivity (**Figure 21** (Right)) versus, e.g. (**Figure 18** (Left)) or (**Figure 19** (Right)).

**Figure 21.**
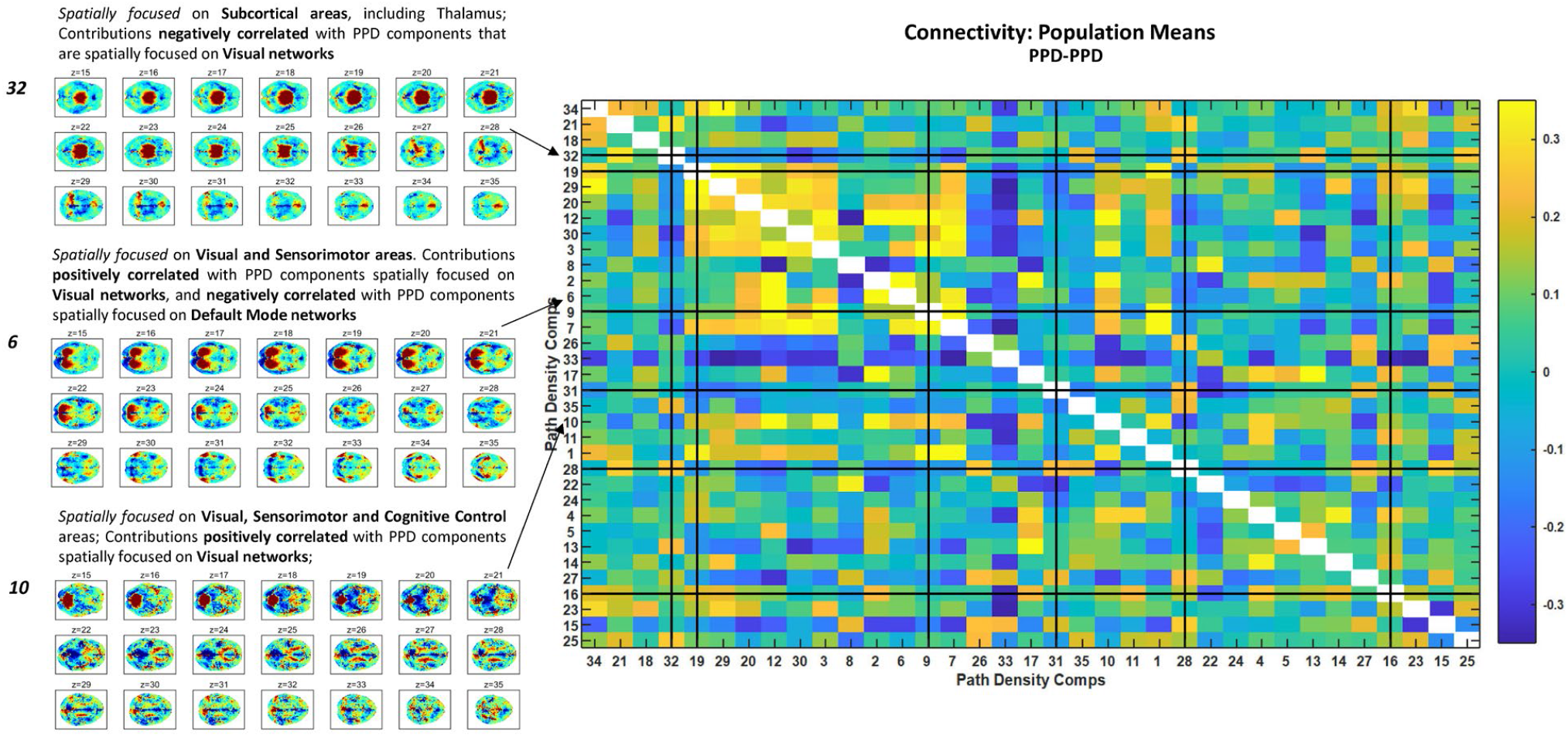
(Rightmost column) Population average spatio-propagative connectivity. The 35 pdGICA components are displayed on both axes, arranged in order of the fnGICA maps to with which they have largest overlap (as in **Figure 16**). The (i,j)^th^ matrix element contains is the average correlation between subject-specific timecourses for the pdGICA timecourses for components *i* and *j*. (Left Column) Select axial slices (z=15 through z=35) of component maps for three of the pdGICA components whose temporal correlations with fnGICA component timecourses are significantly impacted by SZ (see **Figure 20**); pdGICA components 32 is spatially focused in subcortical areas and is negatively correlated with pdGICA components spatially focused on visual networks; pdGICA components 6 and 10 are spatially focused in visual and sensorimotor areas; both are positively correlated with pdGICA coponents focused on visual networks; pdGICA component 6 is weakly negatively correlated with pdGICA components focused on default mode networks while component 10 is weakly positively correlated with pdGICA components focused on default mode networks.

The effects of schizophrenia on spatio-propagative connectivity are relatively sparse but broadly consistent with those observed in functional network connectivity and spatio-functional connectivity. In particular, SZ tends to suppress spatio-propagative connectivity between pdGICA components that are spatially focused in visual areas, and elevates spatio-propagative connectivity between the pdGICA component most spatially focused on thalamus and those that are spatially focused in visual areas (**Figure 22** (Right)).

**Figure 22.**
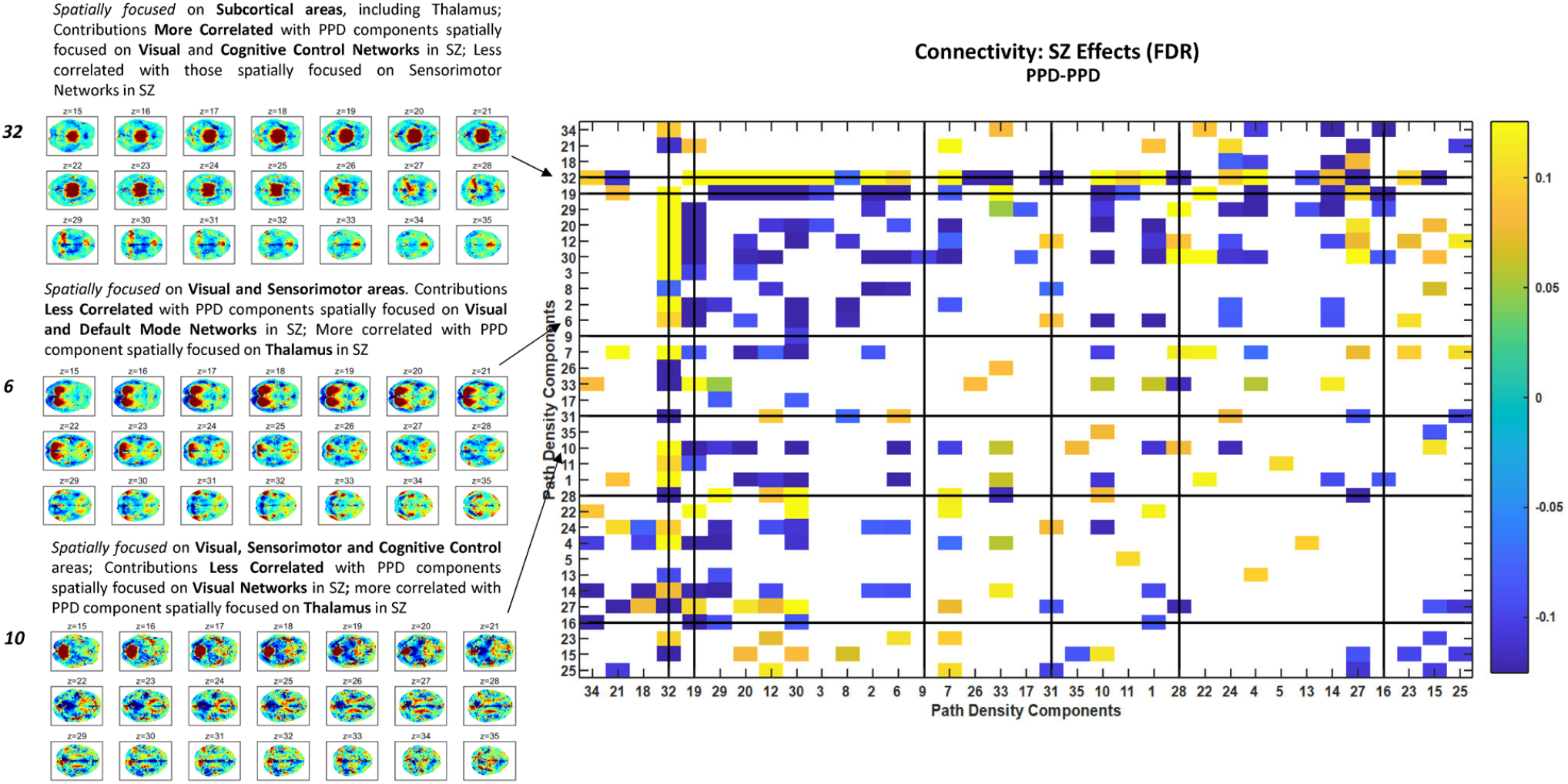
(Rightmost column) SZ effects on spatio-propagative connectivity; (Left Column) Select axial slices (z=15 through z=35) of component maps for three of the pdGICA components whose temporal correlations with fnGICA component timecourses are significantly impacted by SZ (see **Figure 20**); pdGICA component 32 is spatially focused in subcortical areas and is more correlated (or less anticorrelated) among SZ patients with pdGICA components focused on visual and cognitive control networks in SZ patients; pdGICA component 6 is spatially focused on visual and sensorimotor areas and is less correlated in SZ patients with a subset of pdGICA components that are focused on visual, sensorimotor and cognitive control functions; pdGICA component 10 is spatially focused on visual, sensorimotor and cognitive control areas and exhibits lower correlations with many pdGICA components focused on visual areas in SZ patients.

## 4. DISCUSSION

We present here an extensible framework for investigating spatial signal propagation and propagative attractors in BOLD fMRI data based on spatiotemporally localized directional flows in the 4D signal. To our knowledge, highly-localized directional flows in the fMRI signal have not previously been reported upon from a strictly empirical, data-driven perspective. We have exposed spatially structured patterns in, e.g. average direction-specific spatial derivatives (**Figures 9, 10**, 11) that vary significantly between patients with schizophrenia and healthy controls. These local directional signal flows and the broader brainwide propagative consequences present novel, measurable local-global phenomena in the BOLD fMRI signal within which functional and clinical biomarkers might be identified. On a functional level, there is clear regional differentiation in average propagative path density (**Figure 14** (A, B)). Frontal areas are, on average, attracting less spatial signal flow than other regions. In SZ patients, however, these areas have significantly greater propagative density than they do in controls. Schizophrenia also significantly amplifies the tendency of spatial signal flows to concentrate in default mode networks (**Figure 14** (B)). The complex relationships between spatial signal propagation and functional network timeseries has proven difficult to characterize. As expected, there is a positive temporal correlation between propagative path density in a functional network and the network’s contribution to overall signal reconstruction (**Figure 15** (C,D)). Decomposing PPD maps with group ICA produces highly structured component maps that seem to weakly organize along functional axes (**Figure 16**). There are pervasive differences in the importance of these components in reconstructing PPD maps in schizophrenia patients vs. healthy controls (**Figure 17** (B)). The correlation between pdGICA component timeseries and functional network timeseries structure in plausible ways with respect to degree of spatial overlap between the pdGICA component map and the functional network (**Figure 19**). Schizophrenia disrupts correlative relationships between pdGICA component timeseries and functional network timeseries (**Figure 20**). There are certain networks whose correlative relationships with ppdGICA component timeseries are especially disrupted in schizophrenia. These networks include the thalamus, the left posterior central gyrus, the medial temporal gyrus, the inferior frontal gyrus and the anterior cingulate cortex (**Figure 20** (grey columns)). This work is still in its early stages and many limitations remain. An incomplete list of limitations and ongoing/future work would include: better accounting for anatomical artifacts without introducing unnecessary barriers to propagative BOLD signal flow; optimizing the preprocessing pipeline for sensitivity and robustness of this type of analysis; optimizing the post-processing pipeline, e.g. the window length utilized for computing PPD maps, for sensitivity and robustness; modulating the 3mm^3^ discretization of brain space to allow relaxation of currently uniform propagative step size; investigate directed propagation between specific functional regions and time-varying “flow connectivity” via channels of propagative concentration between functional regions. Leveraging the explicit spatial structure in BOLD fMRI to gain better understanding of local-global spatial processes that support or, alternatively, impede healthy functional dynamics could provide more sensitive and condition-specific biomarkers for neuropsychiatric disorders. The extensible framework we introduce here is a preliminary step toward more comprehensively utilizing the local-global spatial and functional information provided by BOLD fMRI.

## Notes

### Competing Interest Statement

The authors have declared no competing interest.

